# GABA/Glutamate neuron differentiation imbalance and increased AKT/mTOR signalling in CNTNAP2^-/-^ cerebral organoids

**DOI:** 10.1101/2024.04.21.590378

**Authors:** Kleanthi Chalkiadaki, Elpida Statoulla, Maria Zafeiri, Georgia Voudouri, Theoklitos Amvrosiadis, Alexandra Typou, Niki Theodoridou, Dimitrios Moschovas, Apostolos Avgeropoulos, Martina Samiotaki, John O. Mason, Christos G. Gkogkas

## Abstract

We developed a human cerebral organoid model derived from induced pluripotent stem cells (iPSCs) with targeted genome editing to abolish protein expression of the Contactin Associated Protein-like 2 (*CNTNAP2)* autism spectrum disorder (ASD) risk gene, mimicking loss-of-function mutations seen in patients. CNTNAP2^-/-^ cerebral organoids displayed accelerated cell cycle, ventricular zone disorganisation and increased cortical folding. Proteomic analysis revealed disruptions in Glutamatergic/GABAergic synaptic pathways and neurodevelopment, highlighting increased protein expression of corticogenesis and neurodevelopment-related genes such as Forkhead box protein G1 (FOXG1) and Paired box 6 (PAX6). Transcriptomic analysis revealed differentially expressed genes (DEG) belonging to inhibitory neuron-related gene networks. Interestingly, there was a weak correlation between the transcriptomic and proteomic data, suggesting nuanced translational control mechanisms. Along these lines we found upregulated Protein Kinase B (Akt)/mechanistic target of rapamycin (mTOR) signalling in CNTNAP2^-/-^ organoids. Spatial transcriptomics analysis of CNTNAP2^-/-^ ventricular-like zones demonstrated pervasive changes in gene expression, particularly in PAX6^-^ cells, implicating upregulation of cell cycle regulation pathways, synaptic and Glutamatergic/GABAergic pathways. We noted a significant overlap of all D30 cerebral organoids ‘omics datasets with an idiopathic ASD (macrocephaly) iPSC-derived telencephalic organoids DEG dataset, where FOXG1 was upregulated. Moreover, we detected increased Glutamate decarboxylase 1 (GAD1) and decreased T-Box Brain Transcription Factor 1 (TBR1) expressing cells, suggesting altered GABAergic/Glutamatergic neuron development. These findings potentially highlight a shared mechanism in the early cortical development of various forms of ASD, further elucidate the role of CNTNAP2 in ASD pathophysiology and cortical development and pave the way for targeted therapies using cerebral organoids as preclinical models.

## Introduction

Elucidating the polygenic basis of neurodevelopmental disorders remains a forefront challenge^1^. ASD is a complex neurodevelopmental condition characterised by challenges in social interaction, communication, and a tendency towards repetitive behaviours and restricted interests^2^. The advent of human iPSC-derived brain organoids, offers a versatile platform to elucidate the molecular and cellular mechanisms implicated in brain development and study genetic and pharmacological models where development can go awry as in ASD, ultimately linking basic research to the development of therapies or to personalised medicine applications^3^.

The *CNTNAP2* gene is one of the largest genes in the human genome, spanning approximately 2.3 million base pairs^4^. CNTNAP2 (or CASPR2) protein belongs to the neurexin family and is a transmembrane protein integral to the function of juxtaparanodes in myelinated neurons^5^. It facilitates the organisation of key proteins like Transient Axonal Glycoprotein 1 (TAG-1) and Potassium Voltage-Gated Channel Subfamily A Member 1 (Kv.1), regulating neuronal excitability^6^. Highly penetrant, homozygous loss-of-function mutations in *CNTNAP2* lead to Cortical Dysplasia Focal Epilepsy (CDFE) syndrome, marked by ASD, intellectual disability, and epilepsy, underscoring its critical role in brain development and function^7^. Importantly, CNTNAP2 has been linked to cortical interneuron development^8^, which in turn have been implicated in the pathophysiology of ASD^9^. Loss of CNTNAP2 results in abnormal neuronal migration and altered distribution of cortical GABAergic interneurons in mouse and zebrafish^8,10^. On the other hand, loss of CNTNAP2 reduced neurite branching and overall neuronal complexity in developing human excitatory neurons^11^. Core phenotypes in *Cntnap2^-/-^* mice were reversed with oxytocin administration^12^. Moreover, *Cntnap2* deletion led to activation of Akt/mTOR signalling in mouse brain^13^. Pharmacological inhibition of this pathway reversed core autism-related phenotypes in *Cntnap2^-/-^* mice^13^. In addition, recent studies reveal that human cortical development is heavily reliant on posttranscriptional regulatory mechanisms, particularly through the regulation by mTOR. mTOR signalling was shown to regulate the architecture of the developing human cortex by maintaining the cytoskeletal organisation in outer radial glial (oRG) cells and the stability of the radial glia scaffold^14^. Furthermore, regulation of translation in early human cerebral organoids progenitor cells, particularly of the 5’Terminal Oligopyrimidine tract (TOR) element-enriched translation machinery, is mediated by mTOR^15^. A recent study using forebrain organoids derived from patient iPSC harbouring *CNTNAP2* c.3709DelG revealed cortical overgrowth and aberrant cellular proliferation because of *CNTNAP2* loss-of-function^16^.

Herein, we employ a human cerebral organoid model derived from a commercially available iPSC line where the CNTNAP2 protein expression was abolished via targeted genome editing, to mimic loss-of-function mutations in ASD patients. CNTNAP2^-/-^ cerebral organoids display altered corticogenesis as evidenced by increased cortical folding, decreased length of cell cycle and disorganisation of ventricular-like neurogenic zones. These phenotypes recapitulate CNTNAP2 organoid models derived from iPSC harbouring loss-of-function mutations^16^. Proteomic analysis of CNTNAP2^-/-^ cerebral organoids revealed disruption of Glutamatergic/GABAergic synaptic pathways and neurodevelopment and increased protein expression of key corticogenesis and neurodevelopmental genes such as FOXG1 and PAX6, in line with accelerated cell cycle and increased cortical folding. Whole transcriptome analysis highlighted significant changes in inhibitory neuron-related gene networks, which were weakly correlated to the proteomics dataset. Spatial transcriptomics in PAX6^+^ cells revealed significant changes in KO versus control, implicating cell cycle regulation pathways in PAX6^+^ cells and on the other hand, synaptic, Glutamatergic/GABAergic pathways in PAX6^-^ cells. We further measured upregulation of Akt/mTOR signalling, in CNTNAP2^-/-^, in line with C*ntnap2^-/-^*mice. Interestingly, we observed a significant overlap of omics datasets to idiopathic ASD (macrocephaly) iPSC-derived telencephalic organoids DEG, where FOXG1 was upregulated, concomitant with overproduction of GABAergic interneurons as evidenced by increased GAD1 expression in CNTNAP2^-/-^, a marker of mature GABAergic cells. Conversely, we detected decreased number of TBR1 expressing cells, a marker of developing glutamatergic layer VI neurons. These data suggest that increased AKT/mTOR signalling in CNTNAP2^-/-^, accompanied by accelerated cell cycle, increased cortical folding, pro interneuron-related transcriptional programmes, and disruption of the Glutamatergic/GABAergic neuron differentiation balance during early cortical development, as shown for other forms of ASD (homozygous PAX6 KO^17^, idiopathic ASD with FOXG1 upregulation^18^), may constitute a shared mechanism during early cortical development of various forms of ASD.

## Results

### CNTNAP2 targeted deletion alters early cortical development in human iPSC-derived cerebral organoids

To study the role of CNTNAP2 in early cortical development, we obtained *CNTNAP2^-/-^* iPSC (KO) generated with Zinc Finger targeted genome editing of both alleles in the XCL1 male human pluripotent stem cell line (control)^19^ (Sup. Fig. 1A). KO and isogenic XCL1 control iPSC displayed normal karyotype (Comparative Genome Hybridisation; CGH array) (Sup. Fig. 1E). and were tested for pluripotency by measuring expression of OCT3/4 and NANOG (with immunofluorescence and RT-qPCR) (Sup. Fig.1F). Using control and KO cells we derived cerebral organoids using a modified protocol (see methods) from refs.^17,20^ (Fig. 1A). Cerebral organoids expressed key typical markers of the early developing brain. Confocal imaging of immunofluorescence labelled cerebral organoid slices at D30 showed prominent expression of the neural progenitor markers SRY-box 2 (SOX2) and PAX6 proteins within the ventricular zone-like structures (VZ, concentric arrangements of columnar epithelial cells around a central cavity, mirroring the structure observed in a neural tube’s cross-sectional view) in both genotypes. Conversely, more mature neurons expressing *β*-Tubulin III (TUJ1) and microtubule-associated protein 2 (MAP2), were located outside the VZ-like structures (Fig. 1B). We detected a 90% reduction in *CNTNAP2* mRNA expression in KO iPSCs and no CNTNAP2 protein expression by immunoblotting in D30 KO organoids compared with control (Sup. Fig. 1C, D). We then proceeded to analyse the morphology of organoids. First, using bright-field microscopy, we observed that D30 KO organoids were ∼8% smaller, compared with control, but no significant size difference was seen at D60, as evidenced by the projected surface area measurements (Fig.1C, D). Second, to assess the proliferation during early development of organoids, we performed cell cycle length assessment by co-labelling with 5-ethynyl-2’-deoxyuridine (EdU) and the proliferating protein marker Ki67 coupled with confocal imaging (Fig. 1E). D30 KO organoids displayed higher EdU/Ki67 ratio (19%), suggesting increased proliferative potential of neural progenitor cells (NPC) and shorter cell cycle duration. (Fig. 1E, F). Third, we proceeded to assess surface folding of cerebral organoids, since expansion and cortical folding in human brain relies on NPC proliferation. We used Surface Electron Microscopy (SEM) to image the gold/palladium-coated surface of D30 cerebral organoids and identified an 82% and 95% increase in surface folding density of D30 and D60 KO organoids, respectively (Fig. 1G, H). Fourth, to further validate the size and the surface folding alterations, we sectioned D30 organoids, and we assessed the size and organisation of VZs, at the histological level. SOX2 positive NPCs and MAP2 positive neurons defined the VZ region boundaries. KO organoids showed significantly increased VZ area (56%) and perimeter (21%) compared with control organoids, while the extensive presence of MAP2 positive cells within the VZ structure indicated a disruption in the organisation of cells (Fig. 1I, J). Disorganisation of VZ cells could impact corticogenesis as shown for other ASD mutations (e.g. Synaptic Ras GTPase-activating protein 1; SYNGAP1 or Phosphatase and tensin homolog; PTEN) modelled with brain organoids^21,22^.

**Figure 1.**
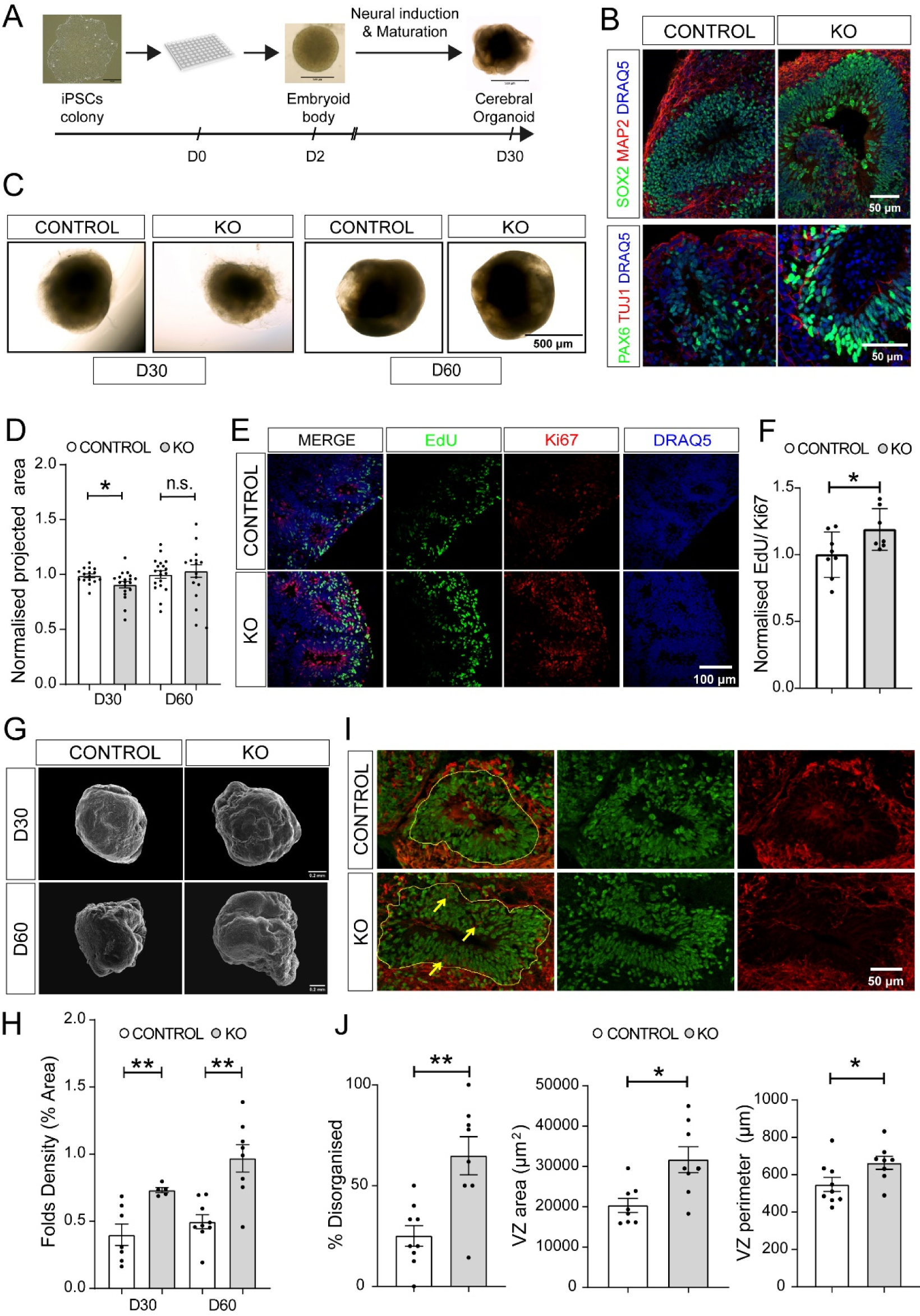
*CNTNAP2* targeted deletion alters early cortical development in iPSC-derived cerebral organoids. **A.** Schematic overview of the cerebral organoid protocol development until day 30 of differentiation. **B.** Representative images from D30 control and KO organoids showing expression of the NPC markers SOX2 and PAX6, and the neuronal markers TUJ1 and MAP2. **C.** Representative bright field images from control and KO organoids. **D.** Normalised projected surface area measurement showing a decrease in size in KO organoids at day 30, which is eliminated at day 60 of differentiation (n=19 for control D30, control D60, KO D30 and n=16 for KO D60, 2 separate organoid batches per genotype for each age group). **E.** Representative images from EdU and Ki67 immunostaining of D30 organoids **F.** Bar graph showing increased EdU/Ki67 ratio in KO organoids (n=7/group, 3 separate organoid batches per genotype, KO values are normalised to control mean). Quantification of cell cycle length was performed using the formula Tc = Ts/ (EdU^+^/Ki67^+^) as previously described ^16^. KO organoids showed shorter cell cycle length compared to control. **G.** Representative scanning electron microscopy images from control and KO organoids. **H.** Quantification of surface fold density showing increased surface folding in KO-derived organoids at day 30 and day 60 of differentiation (n=7 for control D30, n=5 for KO D30, n=9 for control D60 and n=8 for KO D60, 2 separate organoid batches per genotype for each age group). **I.** Representative images from control and KO organoids with SOX2 (green) and MAP2 (red) immunostaining, demonstrating increased size and disorganisation of the VZ in KO organoids. Yellow line delineates the VZ region and yellow arrows emphasize the presence of MAP2 positive neurons inside the VZ. **J.** Quantification of the area and perimeter of VZ and the occurrence of organised versus disorganised VZ regions in control and KO organoids, based on SOX2 and MAP2 staining patterns (n=9 for control and n=8 for KO organoids, 4 separate organoid batches per genotype). For D, F H and J, t test analysis, *p < 0.05, **p < 0.01 See also Sup. Fig. 1 and Sup. Table 7

Altogether, we confirmed accelerated cell cycle in accordance with a previous report employing patient iPSC with loss-of-function mutations^16^. We further revealed increased organoid surface folding and ventricular zone disorganisation, suggesting that the CNTNAP2 KO model used herein affects early cortical development.

### Altered proteomic landscape in CNTNAP2^-/-^ cerebral organoids

To examine the mechanisms underlying *CNTNAP2^-/-^* cerebral organoid phenotypes, we performed label-free phospho-proteomic analysis of D30 CNTNAP2 control and KO cerebral organoids after Matrigel^®^ removal, using the Sp3-mediated protein digestion method^23^, coupled with liquid chromatography–mass spectrometry (LC–MS). DIA mass spectrometry acquisition strategy^24^ enabled the identification of 10,355 protein groups based on 111,568 tryptic peptides (Sup. Table 1). We proceeded with the statistical analysis of the proteome generated based on the proteotypic peptides (peptides unique for a protein), which numbered a total of 8,232 proteins. To test for variation in biological and technical replicates, we performed principal component Analysis (PCA) and confirmed high reproducibility between KO and control samples (Fig. 2B). Proteomic analysis in KO organoids revealed changes in a significant portion of peptides compared with control (-1.5>fold change>1.5; FDR<0.01; *P*<10^-5^; 377 downregulated and 110 upregulated unique peptides) (Fig.2C and Sup. Table 1). Because CNTNAP2 is an autism risk gene we compared this dataset to the SFARI autism gene database and confirmed 31 overlapping genes (Fig. 2C). Among top upregulated targets in KO proteomic analysis, we detected FOXG1, PAX6, Protein Kinase C Beta (PRKCB) and Solute Carrier Family 32 (GABA Vesicular Transporter), Member 1 (SLC32A1) (Fig. 2C). Because FOXG1 was the top upregulated target in KO, we compared the proteomics dataset to a recent RNA sequencing (RNAseq) dataset from human telencephalic organoids derived from idiopathic ASD patient iPSC, where FOXG1 was upregulated^18^ and detected a significant overlap of 163 targets (Fig. 2D). To further understand the pathways downstream of CNTNAP2 deletion in human cerebral organoids, we performed gene ontology (GO) analysis of genes coding for the differentially expressed peptides detected in the proteomics dataset (Sup. Table 2). GO analysis of the 110 upregulated genes showed significant enrichment of terms such as ‘nervous system development’, ‘synapse’, ‘neuron differentiation’, ‘neurogenesis’ and ‘generation of neurons’, while GO terms for the 377 downregulated genes were predominantly related to the extracellular matrix (ECM; organisation, adhesion, collagen) (Fig. 2C, E and Sup. Table 2). Interestingly, upregulated genes in KEGG and SynGO analysis highlighted terms such as: GABAergic and Glutamatergic synapses, pre and post synapses (Fig. 2E, F). GO analysis in the GeDiPNet database (Enrichr) was enriched for terms such as: ‘agenesis of corpus callosum’, ‘autism’, ‘mental retardation’, ‘developmental delay’ and ‘schizophrenia’ (Fig. 2C). Conversely, the downregulated peptides GO dataset analysis showed ECM and focal adhesion pathways and no significant enrichment for synaptic or autism-related pathways (Fig. 2E, F). We also analysed D60 organoids with proteomics (Sup. Fig. 2). We observed changes in protein expression as in D30 (-1.5>fold change>1.5; FDR<0.01; P<10^-5^; 428 downregulated and 156 upregulated unique peptides) (Sup. Fig.2 and Sup. Table 1). GO analysis, akin to D30, revealed upregulation of neurogenesis, synaptic and neuron differentiation pathways and downregulation of ECM-related pathways (Sup. Table. 2).

**Figure 2.**
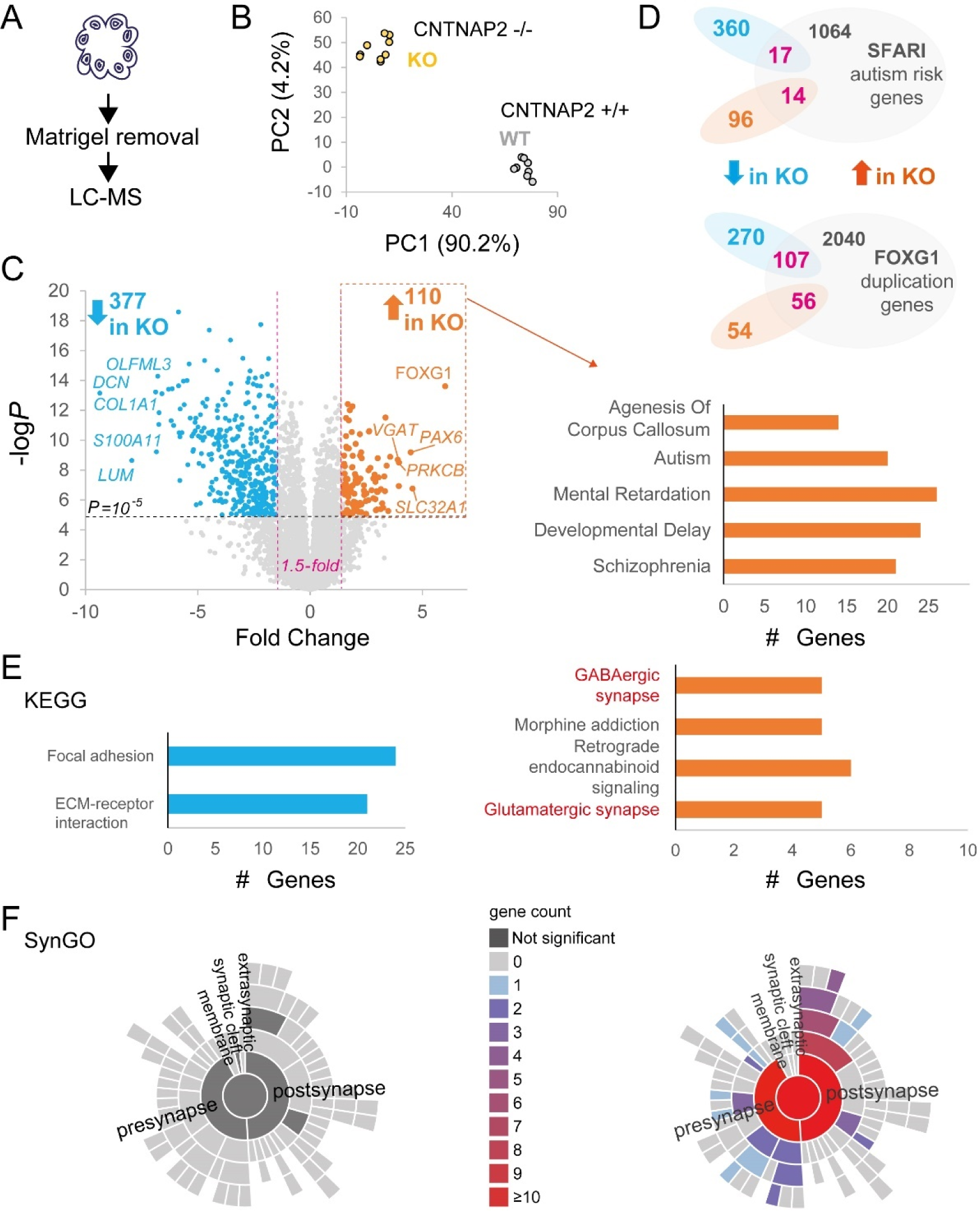
Altered proteomic landscape in CNTNAP2^-/-^ cerebral organoids. **A.** Illustration of cerebral organoid Matrigel^®^ removal and lysis for LC-MS. **B.** Principal Component Analysis (PCA) for proteomics biological (n=3) and technical (n=3) replicates of *CNTNAP2* control (+/+) or KO (-/-) organoids (1 organoid batch per genotype). Scatter plot visualising the proportion of explained variance, with Principal Component 1 (PC1) accounting for the majority (90.2%) of the variance. PC2 explains a smaller portion of the variance (4.2%). **C.** *Left*: Volcano plot of D30 proteomics experiment highlighting upregulated (orange) and downregulated (cyan) peptides in KO samples. X-axis demonstrates the log-transformed fold change in abundance (KO/control) and the Y-axis indicates the log-transformed *P* values associated with individual peptides. A cut-off of ±1.5 fold-change (dashed vertical lines) and *P* value >10^-5^ (dashed horizontal line) was applied. *Right:* Gene Ontology (GO) analysis of upregulated peptides in KO with the GeDiPNet database (Enrichr). **D.** Venn diagrams showing overlap (magenta) of differentially expressed peptides in KO with *top:* SFARI syndromic ASD genes (https://gene.sfari.org/) and *bottom:* RNAseq data from ref.^18^ (FOXG1 upregulation in idiopathic ASD). **E.** KEGG and **F.** SYNGO GO analysis of downregulated (left) and upregulated (right) peptides in KO cerebral organoids. See also Sup. Fig. 2 and Sup. Tables 1 and 2

Taken together, these data show changes in the proteomic landscape of cerebral organoids following CNTNAP2 deletion, revealing upregulation of protein expression linked to neuronal development, autism, and synapses.

### Pro-interneuron transcriptional networks in CNTNAP2^-/-^ cerebral organoids

To further study changes in protein expression at the level of transcription in CNTNAP2^-/-^ cerebral organoids, we performed bulk RNA sequencing of mRNA extracted from D30 cerebral organoids (control or KO) (Fig. 3A). Illumina Novaseq of prepared libraries yielded high quality reads (Sup. Fig. 3). Biological replicates were highly reproducible, as evidenced by the high correlation within genotypes in PCA (Fig 3B). We detected 208 differentially expressed genes (DEG; 42 downregulated and 166 upregulated; (-log *P*adj>1.3, 1.5<log_2_ fold change<-1.5) (Fig. 3C), which overlapped with the SFARI (10 common genes) and FOXG1 upregulation (49 common genes) datasets (Fig. 3D) and is not as extensive as the overlap of the proteomics dataset with SFARI/FOXG1 datasets (Fig. 2D). GO analysis showed enrichment for regulation of transcription, synaptic transmission, axon guidance, neuron fate specification and neuron differentiation (Fig. 3E). Notably, within the top DEGs we detected upregulation of transcription factors associated with cortical interneuron development (such as Distal-Less Homeobox family genes; DLX1, DLX2, NK2 Homeobox 1; NKX2.1, LIM/homeobox protein 6; LHX6), implicated in VZ neurogenesis and cell fate commitment^25,26^ and SVZ (subventricular zone) cell fate commitment and tangential migration^27^ (Fig. 3E).

**Figure 3.**
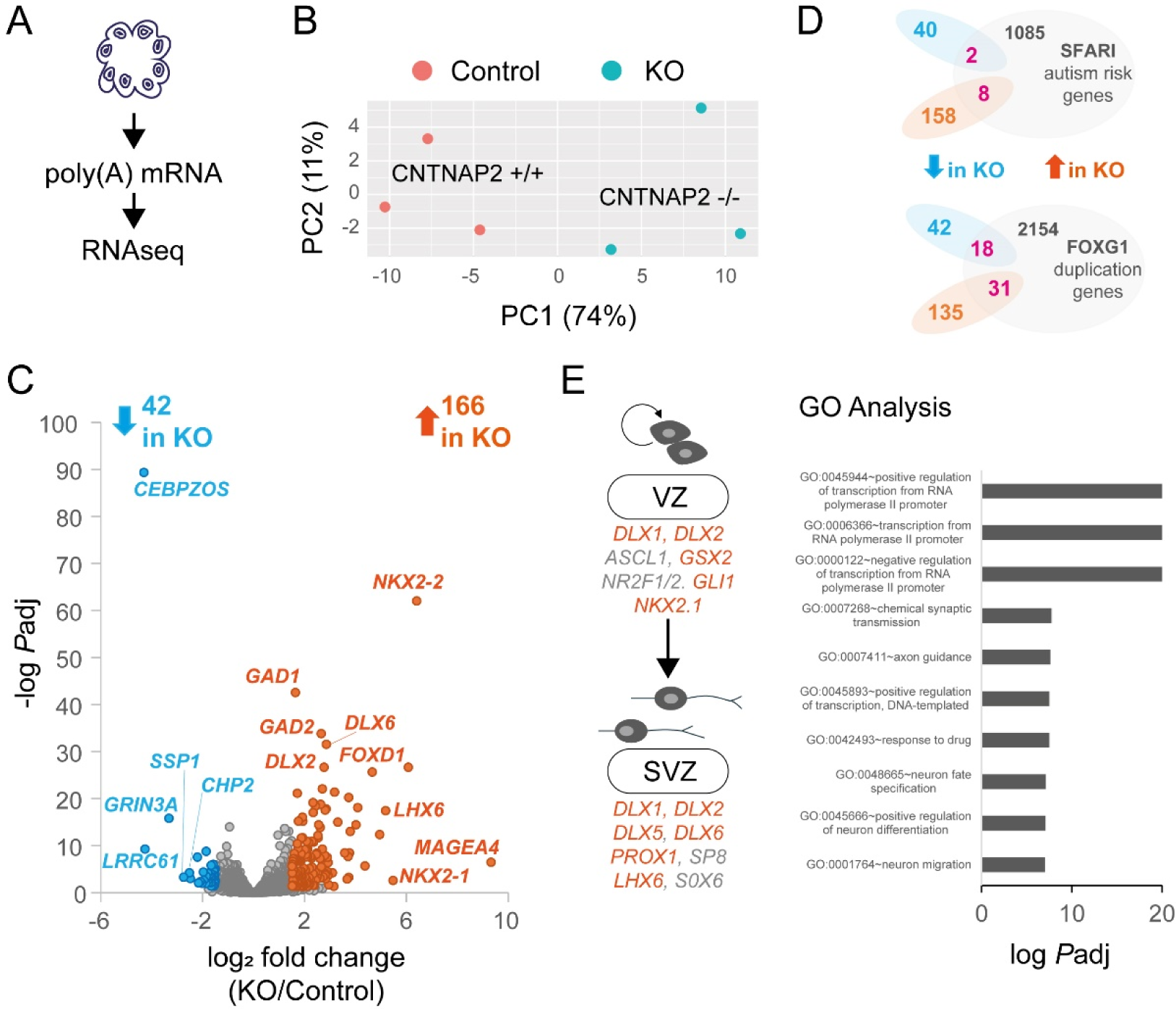
Pro-interneuron transcriptional networks in CNTNAP2^-/-^ cerebral organoids. **A.** Illustration of cerebral organoid poly(A) mRNA isolation and RNAseq. **B.** Principal Component Analysis (PCA) for RNAseq biological (n=3) replicates of *CNTNAP2* control (+/+) or KO (-/-) organoids (1 organoid batch per genotype). Scatter plot visualising the proportion of explained variance, with Principal Component 1 (PC1) accounting for the majority (74%) of the variance. PC2 explains a smaller portion of the variance (11%). **C.** Volcano plot of D30 RNAseq experiment highlighting upregulated (orange) and downregulated (cyan) differentially expressed genes (DEG) in KO samples. X-axis demonstrates the log_2_-transformed fold change in abundance (KO/control) and the Y-axis indicates the negative log-transformed *Padj* (adjusted *P)* values associated with individual mRNAs. A cut-off of ±1.5 fold-change (dashed vertical lines) and *P* value >10^-5^ (dashed horizontal line) was applied. **D.** Venn diagrams showing overlap of DEG in KO with top: SFARI syndromic ASD genes (https://gene.sfari.org/) and bottom: RNAseq data from ref.^18^ (FOXG1 upregulation in idiopathic ASD). **E.** *Left:* Illustration of pro-interneuron mRNAs upregulated (in orange) in CNTNAP2 KO RNAseq DEG, showing genes participating in Ventricular Zone (VZ) and Subventricular Zone (SVZ) neurogenesis, cell-fate commitment, and tangential migration (in grey). *Right:* Gene Ontology (GO) analysis of DEG using GeneSCF v1.1-p2. Significantly enriched GO categories are whine with adjusted *P*-value (padj)<0.05 (Fisher exact test). See also Sup. Fig. 3 and Sup. Tables 1 and 2

Together, these data suggest that *CNTNAP2* loss-of-function may engender pro-cortical interneuron gene expression programmes by upregulating the expression of key VZ and SVZ transcription factors.

### Pervasive changes in gene expression in PAX6^-^ cells in CNTNAP2^-/-^ brain organoids

Given certain phenotypes such as increased cortical folding, accelerated cell cycle and synaptic pathway GO enrichment in D30 KO organoids, we reasoned that *CNTNAP2* deletion could lead to a faster maturation of neuronal progenitors. Moreover, PAX6 protein expression, a marker for neuronal stem cells, was significantly upregulated in KO proteomics analysis (Fig. 2). Thus, we employed spatial transcriptomics analysis using the Nanostring platform GeoMx^®^ (Fig. 4Α) to measure whole transcriptome changes in PAX6^+^ and PAX6^-^ cells, reasoning that as cortical progenitor cells mature, they lose PAX6 expression^28^. D30 cerebral organoid formalin-fixed slices were mounted on glass slides and hybridized with custom-designed barcoded oligonucleotide probes targeting the entire human transcriptome (Fig. 4A). The probes were conjugated with unique molecular barcodes to retain spatial information. Oligos were released (UV illumination) and subjected to bulk RNA sequencing from specific locations (areas of interest; AOI) in the tissue where their corresponding antibodies were bound (Fig. 4Α). To ensure that we captured both cytoplasmic and nuclear RNAs, we used double labelling for PAX6 and NESTIN respectively (Fig. 4Α, Β). Sequencing saturation was >50% and 13,457 genes normalised by 3rd quartile (Q3), expressed above the Limit of Quantitation (LOQ) in at least 1% of AOIs (Sup. Fig. 4A, B, C). Initially, we compared PAX6^-^ and PAX6^+^ cells separately for each genotype (control or KO) (Fig. 4C, D). Whole transcriptome analysis of PAX6^-^/NESTIN^+^ cells versus PAX6^+^/NESTIN^+^ showed overall significantly altered mRNA expression (-log *P*>1.2, 1<log_2_ fold change<-1) which was more prominent in KO (1011 DEG) compared with control (179 DEG) (Fig. 4C). PAX6^-^/NESTIN^+^ cells GO analysis was enriched for synaptic pathways in both genotypes (Fig. 4D), and this effect was more pronounced in KO (537 DEG) compared with control (79 DEG). Notably, we detected 574 DEG in KO PAX6^+^/NESTIN^+^ (compared with PAX6^-^ /NESTIN^+^ cells) with GO enrichment for cell cycle-related pathways, (Fig. 4D), in accordance with the accelerated cell cycle of D30 cerebral organoids (Fig. 1). In contrast, in control organoids PAX6^+^/NESTIN^+^ cells, the 100 DEG were enriched for GO categories such as nervous system development and anatomical structure (Fig. 4D).

**Figure 4.**
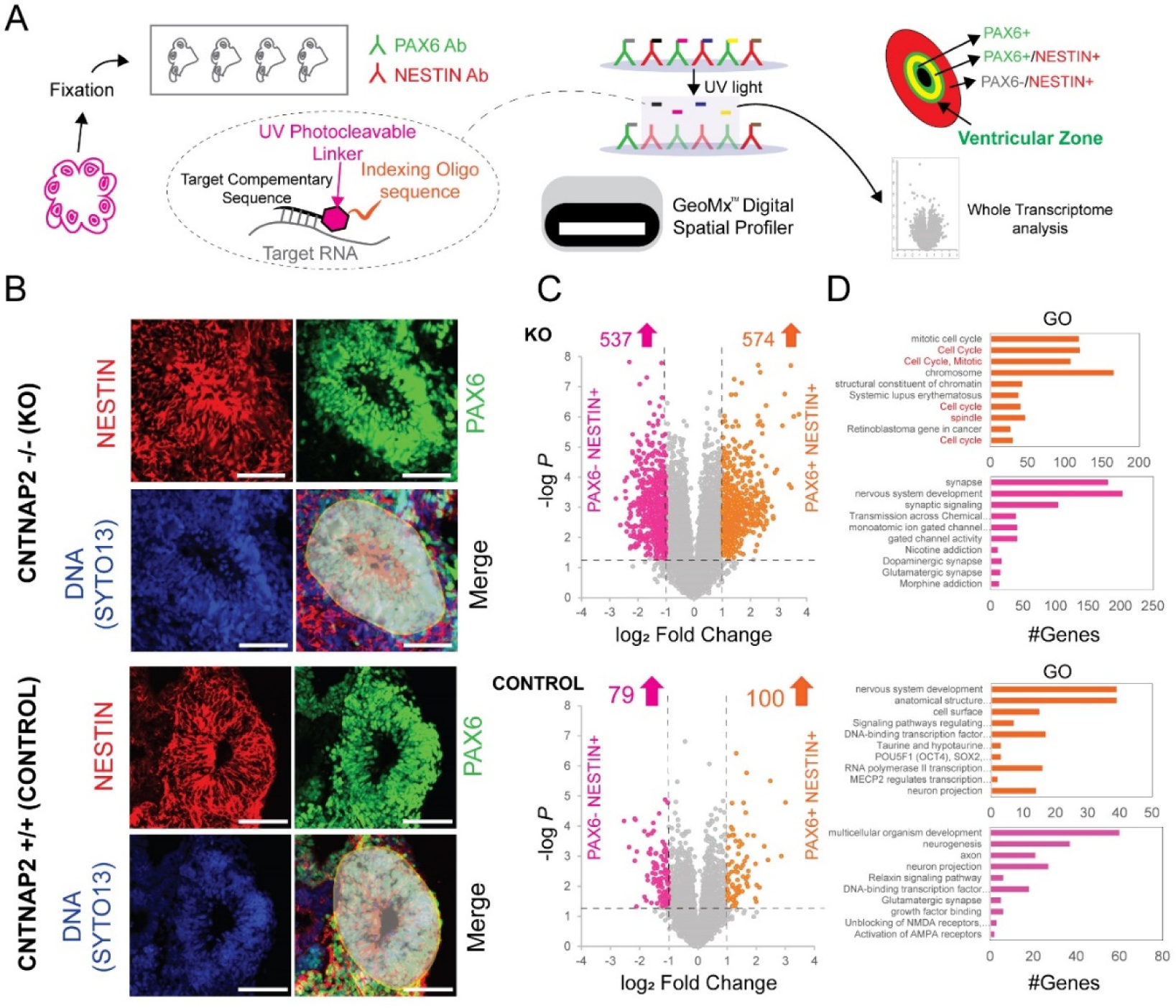
Spatial whole transcriptome analysis (WTA) shows changes in gene expression in PAX6^-^ cells in CNTNAP2^-/-^ brain organoids. **A.** Illustration of experimental procedure to capture PAX6^+^ and NESTIN^+^ WTA with GeoMx^®^ (see text). **B.** Representative images of immunostained slices from D30 control and KO cerebral organoids with GeoMx^®^ wide-field epifluorescence scope. Slices were stained for PAX6, NESTIN and SYTO13 (a nuclear marker labelling DNA). Representative AOIs captured (UV illumination) are shown in grey. C. *Left:* Volcano plot of D30 spatial transcriptomics experiment highlighting upregulated and downregulated differentially expressed genes (DEG) in KO (n=3 biological replicates) or control samples (n=3 biological replicates, 1 batch per genotype) (Pink: PAX6^-^/NESTIN^+^, Orange: PAX6^+^/NESTIN^+^). X-axis demonstrates the log2-transformed fold change in abundance (KO/control) and the Y-axis indicates the negative log-transformed Padj (adjusted P) values associated with individual mRNAs. A cut-off of ±1 log_2_ fold-change (dashed vertical lines) and log *P* value>1.3 (dashed horizontal line) was applied. **D.** Gene ontology analysis of DEG shown in C with g:Profiler. Top categories (Biological Process, Molecular Function and Cellular Compartment) are shown for upregulated and downregulated DEG in KO. Statistical analysis was carried out using g:GOSt (Fisher’s one-tailed test). See also Sup. Fig. 4 and Sup. Tables 5 and 6

Subsequently, we conducted separate gene expression analyses for control and KO cells, distinguishing between PAX6^-^ and PAX6^+^ populations (Sup. Fig. 4E, F). Differential gene expression analysis in PAX6^-^/NESTIN^+^ cells highlighted significantly upregulated synaptic (Glutamatergic/GABAergic), neurodevelopment-related pathways and pre-NOTCH transcription and translation, and downregulated ECM-related pathways in KO compared with control (Sup. Fig. 4E). Notably, in PAX6^+^/NESTIN^+^ cells differential gene expression analysis showed downregulation of ECM-related pathways and upregulation of homeobox (HOX) gene pathways in KO compared with control (Sup. Fig. 4F).

Overall, spatial analysis of gene expression revealed significant changes in CNTNAP2 KO PAX6^+^ cells linked to cell-cycle, in accordance with EdU/Ki67 immunofluorescence and morphological (surface folding) data (Fig.1). Changes in KO transcription versus control were more pronounced in PAX6^-^ compared with PAX6^+^ cells. Along these lines, different transcriptional pathways (NOTCH in PAX6^-^ and HOX in PAX6^+^) were altered in KO compared with control, suggesting that CNTNAP2 may regulate distinct molecular pathways in a cell-type specific manner (PAX6^+^ versus PAX6^-^).

### Weak correlation between Proteomics and RNAseq and altered AKT/mTOR signalling in CNTNAP2^-/-^ brain organoids

Recent studies proposed that posttranscriptional regulatory mechanisms are required for the fidelity of cortical development and that this is largely due to mTOR regulation^14,15^. Along the same lines, PTEN^-/-^ (Phosphatase and Tensin Homolog, upstream of AKT/mTOR) cerebral organoids displayed increased AKT signalling and aberrant cortical development^22^. Moreover, *Cntnap2^-/-^*adult mice displayed increased Akt/mTOR signalling^13^. First, we integrated D30 proteomics and transcriptomics datasets from control and KO cerebral organoids (Fig. 5A). Correlation coefficient of log_2_ fold change in both experiments showed weak inverse correlation (R=-0.2585; Fig. 5Α). Second, we measured AKT/mTOR signalling in cerebral organoids using immunoblotting for key effectors of this pathway: phospho-AKT S473 and phospho-ribosomal protein S6 (rpS6) S240/244 (Fig. 5C). While no significant changes were observed on D30, both p-AKT and p-rpS6 were increased (78% and 30%, respectively) in D60 KO cerebral organoids compared with control (Fig. 5C).

**Figure 5.**
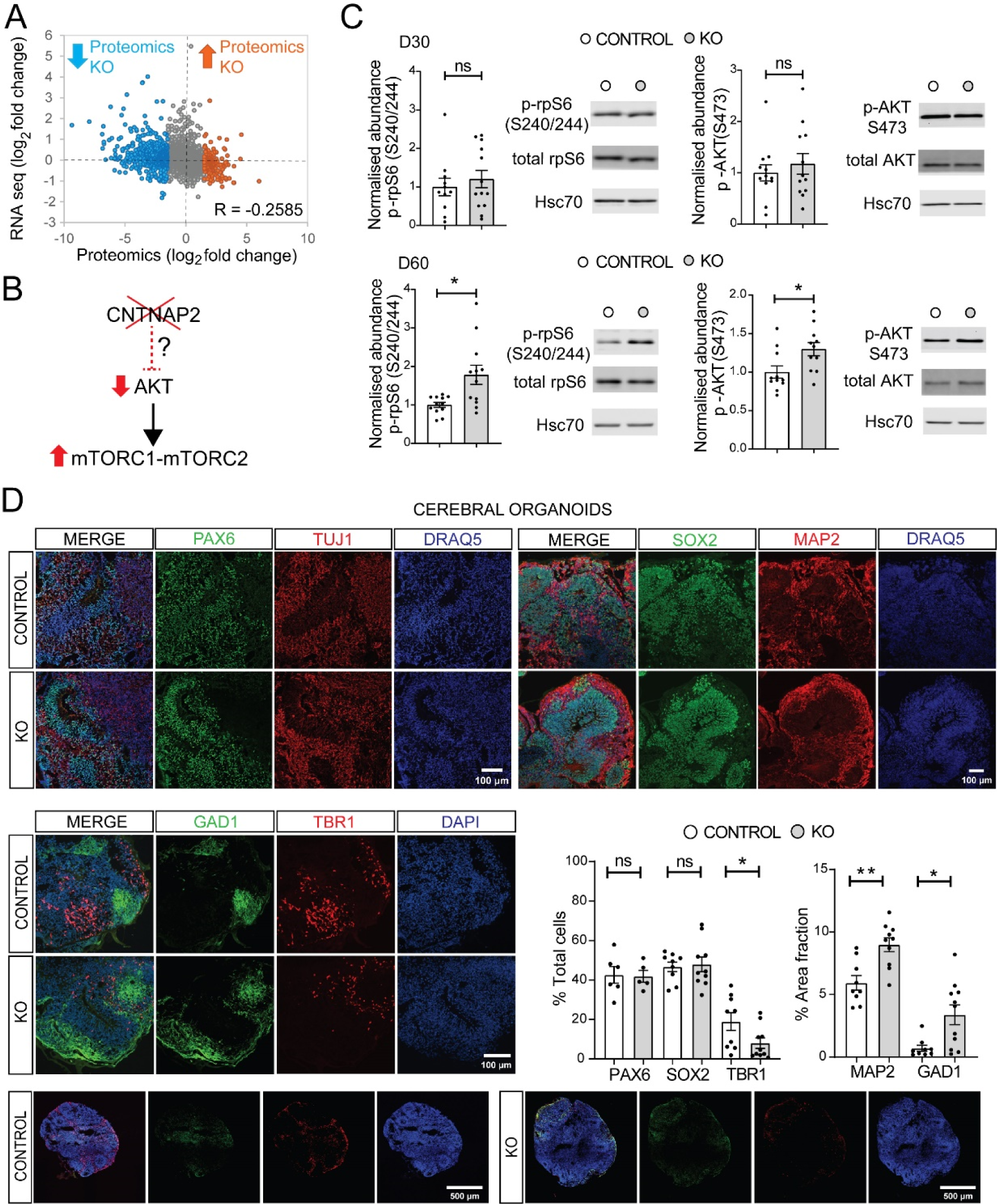
Weak correlation between Proteomics and RNAseq, elevated AKT/mTOR signalling and Glutamatergic/GABAergic fate imbalance in CNTNAP2^-/-^ organoids. **A.** Scatter plot showing Pearson Correlation of Proteomics and RNAseq experiments (R=-0.2585); upregulated and downregulated peptides in KO from the mass spectrometry experiment (Fig. 2) are shown in orange and cyan respectively. **B.** Illustration of possible mechanism for AKT/mTOR activation downstream of *CNTNAP2* loss-of-function **C.** Immunoblot analysis of AKT/mTOR signalling in D30 and D60 KO organoids. *Left:* Quantification of phospho-rpS6 (S240/244) and phospho-AKT (S473) for the indicated groups (n=11-13/group, 4 separate organoid batches for D30, 3 separate organoid batches for D60, KO values are normalised to control mean). *Right*: representative immunoblots of cerebral organoids, probed with antisera against the indicated proteins. HSC70: loading control. **D.** Representative images of immunostaining of SOX2^+^ and PAX6^+^ NPCs, TBR1^+^ postmitotic Glutamatergic neurons and GAD1^+^ mature GABAergic interneurons, depicting the increased MAP2 and GAD1 expression in D30 KO cerebral organoids. *Bottom right:* Quantification of SOX2^+^, PAX6^+^, TBR1^+^ cell fraction, and MAP2 and GAD1 expression (% area fraction) in D30 cerebral organoids. *Bottom:* Representative whole slice images. Total cells were estimated by counting DAPI^+^ or DRAQ5^+^ nuclei (3-6 separate organoid batches per genotype, n=1-3 organoids/batch). For C and D, Student’s *t* test, \*\**P*< 0.01, \**P*<0.01 See also Sup. Fig. 5 and Sup. Table 7

These data suggest that changes in mRNA levels do not consistently correlate with alterations in protein levels in CNTNAP2^-/-^ cerebral organoids, concomitant with upregulation of AKT/mTOR signalling.

### Glutamatergic/GABAergic neuron differentiation imbalance in CNTNAP2^-/-^ brain organoids

Because FOXG1 upregulation was previously linked to Glutamatergic/GABAergic neuron differentiation imbalance in idiopathic ASD patient iPSC-derived telencephalic organoids^18^, and given the enrichment in GO terms linked to Glutamatergic/GABAergic pathways in omics experiments (Fig.2, 3) we reasoned that a similar mechanism may be at play in the CNTNAP2 model. GO analysis in all ‘omics experiments revealed prominent Glutamatergic and GABAergic pathways being regulated (Fig. 2, 3, 4). To confirm this, we performed immunofluorescence analysis and confocal imaging in D30 control and KO cerebral organoids (Fig. 5D) for markers of progenitors (SOX2, PAX6) and of more mature cells (MAP2, TBR1, GAD1). In KO organoids compared with control, we detected increased MAP2 and GAD1 expression, (51% and 380% respectively) (Fig. 5D). Conversely, we measured decreased number of TRB1 positive cells (57%), but no changes in the number of SOX2 or PAX6 positive cells (Fig. 5D), suggesting a shift towards GABAergic neuron progenitors in KO organoids.

Taken together, these data suggest CNTNAP2 KO in cerebral organoids may disrupt the balance of GABAergic/Glutamatergic early neuron differentiation, which to some extent may reflect early maturation of organoids, accelerated cell cycle and increased cortical folding.

## Discussion

The polygenic nature of ASD requires the identification of converging genetic pathways during early development to elucidate its complexity and varied manifestations and propose new therapies^29^. Herein, the cerebral organoid model from iPSCs with targeted genome editing to knock-out CNTNAP2, represents a significant advance in modelling the effects of *CNTNAP2* loss-of-function (Fig. 1), using a commercially available iPSC line with homozygous targeted deletion of CNTNAP2 and an isogenic control line. Previous research utilised patient-derived iPSCs harbouring CNTNAP2 mutations^16^, and the work presented herein offers an additional CNTNAP2 model that can be readily obtained, recapitulating key phenotypes observed in patient-derived organoids. In our model we did not observe significant changes in KO organoid size but report an increase in organoid surface folding (Fig. 1), which is reminiscent of PTEN KO phenotypes^22^. Malformations of cortical folding are common in ASD, epilepsy and cortical focal dysplasias^30^. As in ref.^16^, we further observed accelerated cell cycle and increased FOXG1 and PAX6 expression highlighting the relevance and significance of the new model we developed for studying *CNTNAP2* loss-of-function and ASD. Bulk RNA-seq revealed upregulation of pro-GABAergic interneuron fate transcriptional programmes (Fig. 3). Using spatial transcriptomics (Fig. 4), we further revealed that CNTNAP2 deletion differentially affects PAX6^-^ and PAX6^+^ cells, upregulating cell cycle related DEG in PAX6^+^ cells, and synaptic and Glutamatergic/GABAergic pathways in PAX6^-^ cells. Little is known about the top downregulated DEG in KO: CCAAT/enhancer binding protein zeta opposite strand (CEBPZOS) coding for a mitochondrial protein related to energy metabolism^31^. CEBPZOS downregulation in KO was further confirmed with RNAseq (Fig. 4) and could be linked to GABAergic interneuron cell growth and apoptosis via its mitochondrial localization ^32^. Altogether, these cell-type specific and transcriptional effects further elucidate the role of CNTNAP2 in early cortical development and concur with the early maturation and Glutamatergic/GABAergic neuron imbalance in CNTNAP2 KO organoids (Fig. 5).

We measured significant changes in protein expression (Fig. 2), linked to neurodevelopmental, neurogenic and synaptic pathways, and significant changes in mRNA expression (Fig. 3), boosting pro-interneuron transcription programmes in KO cerebral organoids. The weak correlation observed between transcriptomics and proteomics underscores the complexity of gene expression regulation in cortical progenitors (Fig. 5) and was previously proposed as a new mechanistic explanation for aberrant corticogenesis in ASD^15^. This finding highlights the significant role of post-transcriptional and post-translational mechanisms, particularly through the mTOR pathway, in regulating proteostasis, necessitating deeper investigation.

‘Omics data generated with different methodologies (Figs. 2, 3, 4) in this study showed significant overlap with RNAseq data from telencephalic organoids generated from idiopathic ASD patient iPSC, where FOXG1 was upregulated^18^. FOXG1, a critical transcription factor, plays a key role in the development of cortical interneurons^33^. FOXG1 suppresses the competence to generate the earliest born neurons during later cortical development^34^. FOXG1 conditional deletion has been shown to impair the postnatal distribution of cortical interneurons, leading to enhanced dendritic complexity and decreased migration capacity^35^. Interestingly deletion of *Cntnap2* in mice influences the development and functional integration of interneurons^10^, while CNTNAP2-regulated TAG1 is involved in axonal pathfinding^36^.

Activation of pro-interneuron gene expression following CNTNAP2 deletion, coupled with AKT/mTOR pathway hyperactivation, indicates a potentially key alteration in early neurodevelopmental processes (Fig. 5). This finding suggests that the loss of CNTNAP2 leads to increased expression of transcription factors (such as DLX1/2, NKX2.1 and LHX6), which are known to shape VZ neurogenesis, cell-fate commitment and SVZ tangential migration^25–27^. Zebrafish and mouse *Cntnap2* KO models displayed changes in GABAergic interneuron migration^8,10^. Hyperactivation of the AKT/mTOR pathway downstream of *CNTNAP2* loss-of-function shown herein for a human model, is in accordance with *Cntnap2^-/-^* mouse model results^13^ and could be critical for GABAergic interneuron fate determination and synaptic regulation. Activation of the FOXG1/AKT/Reelin axis was previously shown in focal malformations of cortical development, whereby FOXG1-dependent derepression of Reelin transcription led to misexpression and non-cell autonomous defects in cell migration^37^. Plausibly this aberrant activation may also occur in other forms of syndromic or sporadic ASD where there is AKT/mTOR (PTEN^22^, TSC^15^, SYNGAP1^21^) or FOXG1 (idiopathic ASD^18^) upregulation.

Furthermore, spatial transcriptomics revealed additional transcriptional interplay involving HOX and NOTCH pathways upregulated in CNTNAP2 KO (Fig. 4). HOX genes (HOXD1) map into human chromosome 2 region (2q31-q33), where GAD1 and DLX2 also reside and while linkage to autism is weak for the genes in this region (e.g. GAD1)^38^ the shear proximity could potentially signify transcriptional co-regulation. In parallel, PAX6 homozygous deletion in human iPSC led to upregulation of NOTCH and GABAergic interneuron-related transcriptional programmes (DLX1/2/5/6, GSX2, GAD1/2)^17^. NOTCH affects the proliferation/differentiation balance and DLX, GSX2 govern interneuronal cell fate acquisition. Thus, possibly *CNTNAP2* loss-of-function engenders transcriptional dysregulation, which is critical for cortical proliferation and differentiation and the maintenance of the Glutamatergic/GABAergic progenitor pool balance. Forkhead box protein P2 (FOXP2), the first gene linked to language and speech development^39^ directly binds to regulatory regions of the *CNTNAP2* locus to repress its expression^40^ and was shown to regulate both excitatory and inhibitory neuron development^41^. Interestingly TBR1 was shown to interact with FOXP2 and that TBR1 mutations linked to sporadic ASD disrupt this interaction^42^. Therefore, pro-interneuronal gene expression seen with RNAseq (Fig. 4), in conjunction with the reduced number of TBR1 cells in CNTNAP2 KO organoids (Fig. 5) could be the result of FOXP2-CNTNAP2-TBR1 mediated regulation. Yet, the discordance between transcriptomics and proteomics datasets (Fig. 5) reveals another facet in regulation of gene expression and protein abundance downstream of *CNTNAP2* loss-of-function. This could relate to AKT/mTOR hyperactivation and translational control and operate in parallel or independent to the transcriptional control via CNTNAP2.

In both mouse models^43^ and human studies^44^, CNTNAP2 was linked to the size, structure, or connectivity of the corpus callosum, yet the exact mechanisms of how CNTNAP2 affects the corpus callosum are still unclear. FOXG1 inactivation causes cerebral cortical hypoplasia and corpus callosum hypogenesis^45,46^. Thus, it is conceivable that FOXG1 and CNTNAP2 share developmental roles regarding the proper development of corpus callosum, involving regulation of gene expression via AKT/mTOR or FOXG1-mediated transcriptional programmes.

### Study limitations

We used only one KO iPSC line (generated in XCL1 iPSC; male), which recapitulated most of the cerebral organoid phenotypes previously shown with patient iPSC lines^16^. However, given known issues with human iPSC culture and differentiation variability^47^, it is important to study more clones of the same lines or generate additional (e.g. CRISPR/Cas9 mediated) loss-of-function mutants of *CNTNAP2* in e.g. female iPSC and additional lines, other than XCL1, as those in ref.^16^. For most experiments we used a minimum of 2 separate batches in the analysis, apart from omics, where only 1 batch was used in all experiments. Discrepancies between our model and the patient iPSC-derived organoids^16^ regarding organoid size may reflect iPSC differentiation protocol or imaging methodology differences between the two studies.

## Materials and Methods

### iPSC generation

We used the commercially available iPSC lines XCL1 and XCL1-*CNTNAP2^-/-^*(XCell Science). The bi-allelic CNTNAP2 KO line was generated with the Zinc finger Nuclease (ZFN) method (Sup. Fig. 1Α). An insertion of 4bp and a deletion of 2bp were introduced in allele 1 and 2 of *CNTNAP2* gene in exon 7, respectively. Validation screening for CNTANP2 mutations was performed by Sanger sequencing and pluripotency tests were frequently performed, by measuring the expression of pluripotency markers (Oct3/4 and Nanog) with qPCR and fluorescent ICC (Sup. Fig. 1F). Karyotype analysis was conducted every ∼10 passages using the CGH-array method (Sup. Fig. 1E).

### Cell culture and cerebral organoids generation

iPSC lines were maintained at 37^0^C with 5% CO_2_ in mTSER Plus (STEMCELL Technologies, #05825) on Matrigel^®^ (Corning, #354277) coated plates and passaged using 0.5mM EDTA (Thermo Fisher Scientific, #15575020). Cerebral organoids were generated using a modified organoid differentiation protocol^17^. Briefly, 80% confluent iPSC colonies were dissociated into single cells with Accutase (Sigma-Aldrich, #A6964). A total of 9,000 cells were plated in each well of ultra-low attachment 96-well U-bottom plate (Corning, #7007). Embryoid bodies (EBs) were formed within the next 24 hours (day 1) and kept in the 96 well-plate for 6 days. EB medium changes were performed on day 2 and day 4. On day 6, EBs were transferred to ultra-low attachment 24-well plates (Corning, #3473) and the medium was switched to neural induction medium (Lancaster 2013), with the addition of 10μM SB431542 (Selleck Chemicals, #S1067) and 100nM LDN-193189 (Selleck Chemicals, #S2618). On day 10 EBs were embedded on Matrigel GFR^®^ (Corning, #354230) droplets and cultured in cerebral organoid differentiation medium. Embedded EBs were kept on stationary culture for 24hrs, followed by transfer to an orbital shaker (Heathrow Scientific, #5003396). In order to aid neuronal maturation, 20ng/ml BDNF (Peprotech, #450-02) and NT3 (Peprotech, #450-03) was added to the medium at D30, for two weeks^17^. Organoids were maintained on shakers for 60 days and half medium changes were performed every other day.

### Proteomics sample preparation using the Sp3-mediated protein digestion protocol

D30-D33 organoids were transferred from culture plates, briefly washed in ice-cold DPBS (PAN-Biotech, #P04-36500), depleted of Matrigel using the cell recovery solution (Corning #354253), supplemented with protease and phosphatase inhibitors, and subsequently lysed in a lysis buffer consisting of 4% SDS, 0.1M DTT, 0.1M Tris pH 7.4. For each biological replicate, 3-4 organoids were pooled and lysed together. The samples were sonicated (Digital Sonifier 250 Marshall Scientific), heated for 3 min at 95^0^C, followed by a centrifugation step for 15 min at 17000 x g. The lysed samples were processed according to the Sp3 protocol^23^ including an alkylation step in 200 mM iodoacetamide (Acros Organics). 20 ug of beads (1:1 mixture of hydrophilic and hydrophobic SeraMag carboxylate-modified beads, GE Life Sciences) were added to each sample in 50% ethanol. Protein clean-up was performed on a magnetic rack. The beads were washed two times with 80% ethanol and once with 100% acetonitrile (Fisher Chemical). Proteins captured on beads were digested overnight at 37^0^C under vigorous shaking (1200 rpm, Eppendorf Thermomixer, #5382) supplemented with 0.5 µg Trypsin/LysC (MS grade, Promega) prepared in 25 mM ammonium bicarbonate. Next day, the supernatants were collected, and the peptides were purified using a modified Sp3 clean up protocol and finally solubilized in the mobile phase A (0.1% formic acid in water), sonicated and the peptide concentration was determined through absorbance at 280 nm measurement using a Nanodrop instrument (ThermoFisher, Nanodrop One C).

### LC-MS/MS

Samples were processed on a liquid chromatography tandem mass spectrometry (LC-MS/MS) setup consisting of a Dionex Ultimate 3000RSLC online with a Thermo Q Exactive HF-X Orbitrap mass spectrometer. Peptide samples were directly injected and separated on an 25 cm-long analytical C18 column (PepSep, 1.9μm3 beads, 75 µm ID) using a 90 minutes long run, starting with a gradient of 7% Buffer B (0.1% Formic acid in 80% Acetonitrile) to 28% Buffer B for 63 min, followed by an increase to 36% in 7 min and a second increase to 95% in 0.5min, and, then kept constant for equilibration at 7% Buffer B for 14.5min. A full MS was acquired in profile mode using a Q Exactive HF-X Hybrid Quadrupole-Orbitrap mass spectrometer, operating in the scan range of 375-1400 m/z using 120K resolving power with an AGC of 3x 106 and max IT of 60ms followed by data independent analysis using 8 Th windows (39 loop counts) with 15K resolving power with an AGC of 3x 105 and max IT of 22ms and a normalised collision energy (NCE) of 26. Each biological sample was analysed in three technical replicates.

### LC-MS data analysis

Orbitrap raw data was analysed in DIA-NN 1.8 (Data-Independent Acquisition by Neural Networks)^24^ through searching against the reviewed Human Uniprot database containing 27246 proteins (retrieved 4/2021) in the library free mode of the software, allowing up to two tryptic missed cleavages. A spectral library was created from the DIA runs and used to reanalyse them. DIA-NN default settings have been used with oxidation of methionine residues and acetylation of the protein N-termini set as variable modifications and carbamidomethylation of cysteine residues as fixed modification. N-terminal methionine excision was also enabled. The match between runs (MBR) feature was used for all analyses and the output (precursor) was filtered at 0.01 FDR and finally the protein inference was performed on the level of genes using only proteotypic peptides. The generated results were processed statistically and visualised in the Perseus software (1.6.15.0)^48^. Values were log_2_ transformed, a threshold of 70% of valid values in at least one group was applied and the missing values were replaced from normal distribution. For statistical analysis, Student’s t-test was performed, and permutation-based FDR was calculated. Principal Component Analysis (PCA) was performed using Perseus standard settings.

### Immunofluorescence and confocal imaging

D30 organoids were fixed for 1hr in 4% (w/v) paraformaldehyde in PBS at room temperature (RT), cryo-protected in 30% (w/v) sucrose in PBS, O/N, or until they sank, at 4^0^C, and embedded in optimal cutting temperature (O.C.T.) compound (Shakura Finetek USA Inc, #4583). O.C.T. embedded organoids were frozen using a dry ice/ethanol bath and stored at -80^0^C until further processing. 15μm-thick organoid sections were obtained in a cryostat (Leica cryostat), blocked in 10% (v/v) normal goat-serum (NGS) solution, containing 0.3% TritonX-100/PBS, and incubated O/N in primary antibodies, at 4^0^C. All primary antibodies were detected using AlexaFluor secondary antibodies (Invitrogen), incubated for 90min at RT. A detailed list of antibodies can be found in Sup. Table 8. 4′,6-diamidino-2-phenylindole; DAPI (Abcam, #ab228549) and DRAQ5 (Abcam, ab108410) were used for nuclear staining. Imaging was performed on a LEICA SP5 and a NIKON A1R HD confocal microscope, using a X40 oil-immersion objective, a X20 air objective or a X20 oil-immersion objective (whole slice imaging). For all immunofluorescence experiments a minimum of 3 slices/ organoid were imaged and analysed.

### EdU click-it assay

D30 organoids were incubated in 10μM EdU (Invitrogen, #C-10337, Component A) diluted in cerebral organoid differentiation medium, for 2hrs and subsequently collected and processed as described previously, for immunofluorescence experiments. EdU click-it assay was performed in organoid cryosections per manufacturer’s instructions (EdU kit Invitrogen, #C-10337) followed by immuno-staining for Ki67. The Edu+/Ki67+ ratio was measured, and cell cycle length was calculated using the formula Tc=Ts/(EdU^+^/Ki67^+^) (Tc=cell cycle length, Ts= S phase length). A minimum of 3 slices/ organoid were imaged and analysed.

### Immunoblotting

D30 or D60 organoids were transferred from culture plates, briefly washed in ice-cold DPBS (PAN-Biotech, #P04-36500), depleted from Matrigel^®^ using the cell recovery solution and subsequently homogenized in RIPA buffer as in ref.^49^, supplemented with protease and phosphatase inhibitors, using a motorised pestle mixer. For each biological replicate, 3-5 organoids were pooled and lysed together. Samples were incubated on ice for 15 min, with occasional vortexing and centrifuged for 20 min at 16 000 g at 4°C. Protein concentration of each sample was determined using the BCA protein assay (Pierce™ BCA Protein Assay, Thermofisher). 40 micrograms of protein per lane were prepared in SDS sample buffer (50 mM Tris pH 6.8, 100 mM DTT, 2% SDS, 10% glycerol, 0.1% bromophenol blue), heated to 95^0^C for 5min and resolved on polyacrylamide gels. Proteins were transferred to 0.2 μm nitrocellulose membranes (Bio-Rad), blocked for 1 hr at RT in 5% bovine serum albumin (BSA) in TBS-T and incubated with primary antibodies O/N at 4^0^C. Fluorescent secondary antibodies were used for all immunoblotting experiments. A detailed list of antibodies can be found in Sup. Table 8. Blots were imaged using an Azure imaging system (Azure Biosystems) and quantified using the Image Studio Software (Li-COR Biosciences), by measuring the intensity of each protein band. HSC70 or β-Actin was used as loading control. Data are shown as arbitrary units (AU) as a proxy for protein expression, after normalisation to control (for protein phosphorylation: phospho-protein values were divided to normalised total protein values, after subtracting immuno-blot background intensity). For each experiment, values from KO organoids were normalised to the mean of the control group.

### RNA sequencing and Bioinformatics Analysis

D30 organoids were transferred from culture plates to 1.5ml tubes and briefly washed in ice-cold DPBS. For each biological replicate, 3-4 organoids were pooled in the same tube. After complete removal of DPBS, organoids were homogenized using QIAshredder homogenizers (Qiagen, #79656) and total RNA was extracted using the RNeasy Micro kit (#74004, Qiagen) per manufacturer’s instructions. RNA was suspended in RNase-free water and the concentration and purity of the samples was determined using a Nanodrop instrument (ThermoFisher, Nanodrop One C). Library preparation and RNA sequencing were performed as a service by GENEWIZ/AZENTA and sequenced on a Novaseq 6000 instrument (Illumina).

Sequence reads were trimmed to remove possible adapter sequences and nucleotides with poor quality using Trimmomatic v.0.36. The trimmed reads were mapped to the Homo sapiens GRCh38 reference genome available on ENSEMBL using the STAR aligner v.2.5.2b. BAM files were generated in this step. Unique gene hit counts were calculated by using featureCounts from the Subread package v.1.5.2. The hit counts were summarised and reported using the gene_id feature in the annotation file. Only unique reads that fell within exon regions were counted. After extraction of gene hit counts, the gene hit counts table was used for downstream differential expression analysis. Using DESeq2, a comparison of gene expression between the KO-control groups of samples was performed. The Wald test was used to generate p-values and log2 fold changes. Genes with an adjusted p-value<0.05 and absolute log2 fold change>1 were called as differentially expressed genes for each comparison. A gene ontology analysis was performed on the statistically significant set of genes by implementing the software GeneSCF v.1.1-p2. The goa_human GO list was used to cluster the set of genes based on their biological processes and determine their statistical significance.

### Spatial transcriptomics with GeoMX^®^ and Bioinformatics Analysis

Spatial transcriptomics was performed using the nanoString GeoMx^®^ Digital Spatial Profiler in D30 cerebral organoids. Tissue preparation. D30 organoids were transferred from culture plates to pre-chilled DPBS, washed quickly twice and fixed in 10% (v/v) neutral buffered formalin (NBF), at RT for 24 hrs with gentle agitation. On day 2, organoids were washed twice for 30min in DPBS, followed by sequential incubations in sucrose solutions (10%, 20% and 30% w/v), prepared in RNAse free water, on ice for 1-2 hrs or until tissue sank. Organoids were then incubated in 50:50 OCT-30% sucrose solution for 30min on ice, before O.C.T. embedding and freezing in dry ice/ethanol bath. 10μm-thick organoid sections were obtained in a cryostat and stored at -80^0^C until further processing. Sections processing. Fixed frozen slides were equilibrated to RT, washed for 5min in DPBS to remove O.C.T. and baked for 30min at 60^0^C, in order to prevent tissue detachment from the slides. Slices were then dehydrated in serial incubations in ethanol solutions (50% - 100% v/v), air dried for 15-30min, before rehydration and target retrieval step. Slides were incubated in a preheated Tris-EDTA solution (eBioscience™ IHC Antigen Retrieval Solution - High pH, #00-4956-58) for 15min, at 85^0^C, followed by proteinase K treatment (at a concentration of 0.1 μg/mL for 15 minutes at 37^0^C). In situ hybridization. Overnight hybridization at 37°C was performed using hybridization probes from the Human Whole Transcriptome Atlas (WTA). The slides then underwent two 5min washes, combining equal parts of 4× SSC buffer and formamide. Post-washing, slides were blocked for half an hour and antibodies were applied to identify morphological markers, assisting in the delineation of specific areas of interest (AOIs). We used PAX6 (Cell Signaling Technology, #60433S), NESTIN (Cell Signaling Technology, #33475S) and SYTO83 (GeoMx antibody panel) antibodies as our markers. Following O/N incubation at 4^0^C, slides were washed 4 times in 2x SSC buffer and incubated with secondary antibodies and nuclei marker Syto83, in Buffer W for 1 hour at room temperature in a humidified chamber. Slides were washed 4 times in 2x SSC buffer and placed in the nanoString GeoMx^®^ DSP instrument.

UV light was targeted onto each AOI to facilitate the release of RNA-ID carrying oligonucleotides from the WTA probes, allowing for their individual collection in distinct wells for every AOI. Subsequently libraries were prepared adding Illumina i5 and i7 dual indexing primers to the oligonucleotide tags during PCR for unique indexing of each AOI. AMPure XP beads (Beckman Coulter) were used for PCR cleanup, and library concentration was determined using a Qubit 3.0 fluorometer (Thermo Fisher Scientific). The quality of libraries was assessed with a Bioanalyzer (Agilent), and sequencing was performed on an Illumina NovaSeq 6000 system. Gene count data for each AOI was derived from raw .fastq files using the GeoMx^®^ NGS Pipeline. Data analysis was conducted using the GeoMx DSP online analysis platform (Nanostring). Raw reads were trimmed, stitched, aligned, and deduplicated, using standard settings. Differential expression analysis, focusing on the comparisons between PAX6^-^NESTIN^+^ and PAX6^+^NESTIN^+^ in both control and KO and vice versa, KO versus control in PAX6^-^NESTIN^+^ and PAX6^+^NESTIN^+^ cells. Permutation q-values (*P*-adjusted) were calculated for robust assessment of statistical significance. DEGs were calculated using a cut-off of ±1 log_2_ fold-change and log *P* value >1.3. Principal Component Analysis was performed using the Q3 normalised read counts output by the online GeoMx DSP analysis platform. Gene Ontology (GO) analysis was performed with g:Profiler. Top GO categories (Biological Process, Molecular Function and Cellular Compartment) for DEG were ascertained with statistical analysis using g:GOSt (Fisher’s one-tailed test).

### Scanning Electron Microscopy (SEM)

Scanning electron microscopy (SEM) images were obtained using a JEOL JSM-6510 LV SEM Microscope (JEOL Ltd., Tokyo, Japan) equipped with an X–Act EDS-detector by Oxford Instruments, Abingdon, Oxfordshire, UK (an acceleration voltage of 20 kV was applied). Prior to SEM analysis, the samples (control and KO organoids) were coated with an Au/Pd thin film (4–8 nm) in a sputtering equipment (SC7620, Quorum Technologies, Lewes, UK).

### RT-qPCR

Total RNA was extracted from control and KO iPSCs using TRI reagent (Sigma-Aldrich, #T9424) per manufacturer’s instructions. RT-qPCR was performed using the LUNA 1-step RT-qPCR kit (E3005L, NEB) on an AriaMx Real-time PCR system (G8830A, Agilent Tech.). Raw Ct values were normalised to GAPDH using the ΔΔCt method^50^. Primer sequences (F: forward, R: reverse) used in this study; *OCT4:* F: 5’-GGAGGAAGCTGACAACAATGAAA-3΄, R: 5’- GGCCTGCACGAGGGTTT-3΄, *NANOG*: F: 5΄-ACAACTGGCCGAAGAATAGCA-3΄, R: 5΄- GGTTCCCAGTCGGGTTCAC-3΄, *CNTNAP2 RT-qPCR:* F: 5΄-TGCCTAGAGAGATACCACGGTTACT- 3΄, R: 5΄-TTATATCGTAGCCACATCCCTTCTT-3΄, *CNTNAP2 Genotyping 1*: 5΄- TCTTCCATTGATTTTGCCATCGACC-3΄, R: 5΄-CAGGCTCTTAAAAATCAACAGAGGGAAGC-3΄, *CNTNAP2 Genotyping 2*: F: 5΄-TTTGTAGGACGTGACAGGCTTAGATG-3΄, R: 5΄- AGGGAAGCCATGGTTACCTTTCC-3΄, *GAPDH*: F: 5΄-ACCACAGTCCATGCCATCAC-3΄, R: 5΄- TCCACCACCCTGTTGCTGTA-3΄.

### Image analysis

Bright field images were acquired on an EVOS^TM^ XL Core microscope (Invitrogen). Quantification of projected surface area of the organoids was performed using Image J software, by outlining the area of the organoid. Fold density measurement in SEM images was performed as previously described ^22^ using the canny edge detection plug-in Image J. In fluorescent images, cell counting was performed either manually (for EdU assay experiment), using the cell counter tool in Image J or according to ref.^51^ instructions for cell counting (SOX2, PAX6, TBR1, DAPI and DRQ5). The cell fraction for each marker was calculated as a % of the total nuclei measured. For SOX2^+^ and PAX6^+^ cell fraction, whole images (20X objective) containing VZ structures were measured and analysed. For MAP2 and GAD1 fluorescent staining measurements, whole slice images were obtained and the MAP2 or GAD1 area fraction of each slice was calculated^51^. For the VZ analysis, SOX2 and MAP2 staining was used to manually define the VZ boundaries. Area and perimeter of each VZ was measured using Image J (Analyse/Measure tool). Disorganisation of the VZ was calculated using MAP2 staining as an indicator; a VZ containing MAP2^+^ cells was scored as disorganised, while absence of MAP2^+^ cells (clear boundaries) assigned a VZ as organised. For each organoid a minimum of 5 VZ from at least 3 separate slices were measured and analysed.

### *Cntnap2* KO mice brain tissue collection

3–4-month-old *Cntnap2^-/-^* male mice were obtained from JAX laboratories (Strain #017482). All procedures were in accordance with UK Home Office and University of Edinburgh regulations. Animals were backcrossed for more than 10 generations to C57Bl/6J background. Food and water were provided ad libitum. Weaning of pups was carried out at postnatal Day 21. Mice were then group housed (4-5 littermates per cage) by sex and genotype. Cages were maintained in ventilated racks in temperature (20–21°C) and humidity (∼55%) controlled rooms, on a 12-h circadian cycle (7 a.m.–7 p.m. light period). For brain tissue collection, animals were anaesthetized with isoflurane and culled. Brain tissue was rapidly extracted, washed thrice in ice cold PBS, and lysed for immunoblotting in RIPA buffer using a Dounce glass homogeniser.

### Statistical Analysis

Experimenters were blinded to genotype during image analysis. All data are presented as mean ± S.E.M. (error bars). No randomisation was performed in this study. Normality was assessed using the Shapiro-Wilk test (α = 0.05) and statistical significance was set a priori at 0.05 (n.s.: non-significant). N number corresponds to biological replicates (single organoid, or pooled organoids), unless otherwise specified. If multiple observations per research object were collected, we performed nested data statistical analysis (specified in figure legends and Sup. Table 7). Details for statistical tests used are provided within figure legends or the relative methods description and summarized in Sup. Table 7. Sup. Fig. 8 contains raw immunoblot data. Statistical analysis was performed using GraphPad Prism 9.

## Author Contributions

Conceptualisation, K.C., C.G.G.; Methodology, all authors; Investigation/Methodology, all authors; Writing – Original Draft, K.C., J.O.M and C.G.G. Writing – Review & Editing, all authors; Funding Acquisition, C.G.G.; Supervision, K.C., J.O.M. and C.G.G.; D.M. and A.A. performed SEM experiments.

## Acknowledgments

The authors wish to thank M. Barbato and C. Chan for technical support with brain organoids. We also thank Nanostring for access to the GeoMx^®^ DSP through the Technology Access Program (TAP) and GENEWIZ/AZENTA for RNAseq services.

## Data Availability

The mass spectrometry proteomics data will be deposited to the ProteomeXchange Consortium via the PRIDE^52^ partner repository. RNAseq and spatial transcriptomics (GeoMx^®^) raw data will be deposited to Mendeley. Raw immunoblot images are provided in Sup. Fig. 5. All raw data for this study are available upon reasonable request to the corresponding author.

