## Supplementary Figures and Legends for "GABA/Glutamate neuron differentiation imbalance and increased AKT/mTOR signalling in CNTNAP2^-/-^ cerebral organoids"

**Supplementary Figures 1-5**

**Supplementary Figure 5** Uncropped immunoblot images.

**Supplementary Table 1** Differentially expressed peptides in D30 and D60 cerebral organoids proteomics.

**Supplementary Table 2** Gene Ontology analysis in D30 and D60 cerebral organoids proteomics.

**Supplementary Table 3** Differentially expressed genes in D30 cerebral organoids RNAseq.

**Supplementary Table 4** Gene Ontology analysis in D30 cerebral organoids RNAseq.

**Supplementary Table 5** Differentially expressed genes in D30 cerebral organoids spatial transcriptomics.

**Supplementary Table 6** Gene Ontology analysis in D30 cerebral organoids spatial transcriptomics.

**Supplementary Table 7** Statistical Analysis Details.

**Supplementary Table 8** Antibodies used in this study.

### Supplementary Fig. 1

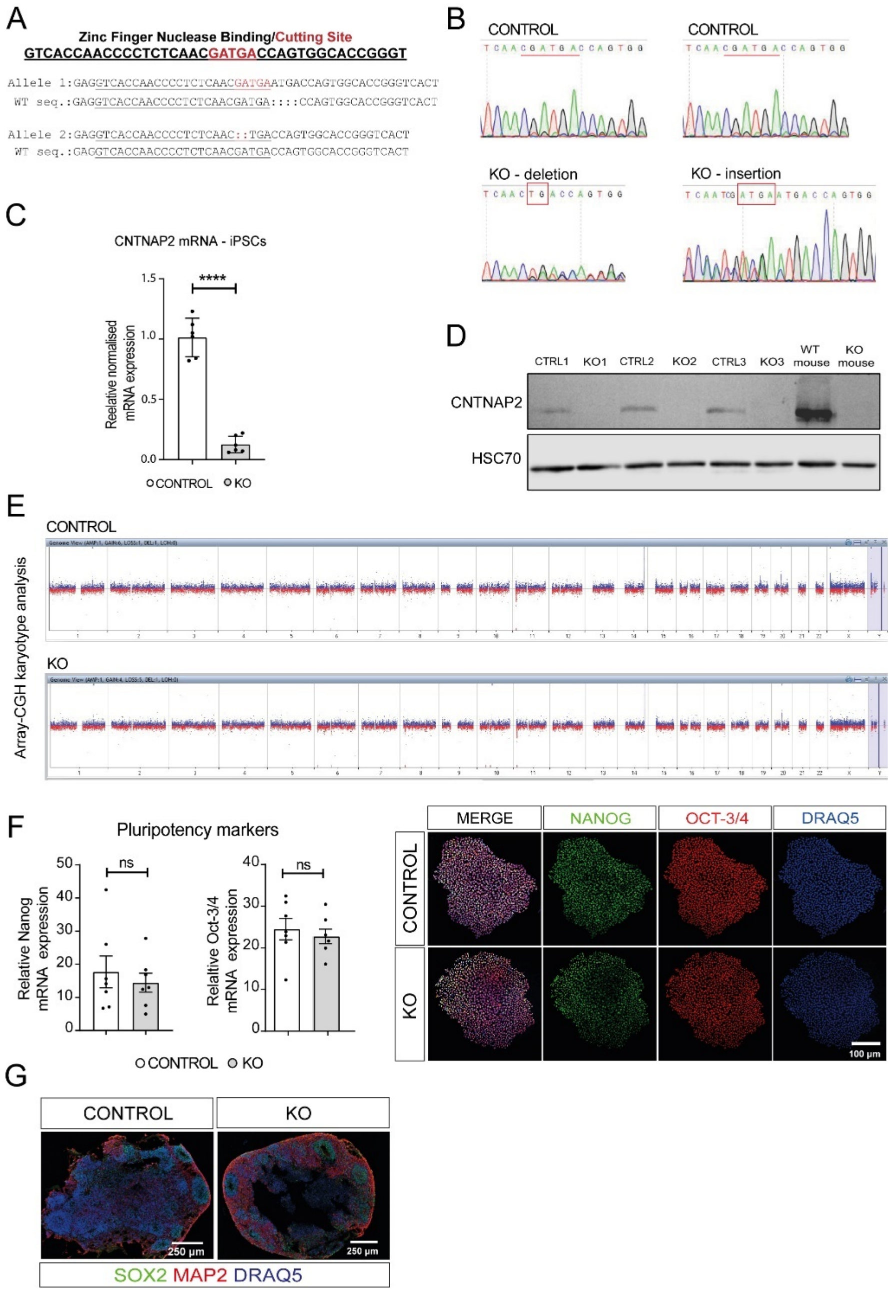


**Supplementary Figure 1 IPSC QC and additional data**. **A.** Bi-allelic CNTNAP2 KO iPSCs generation with the insertion of 4bp (allele1) and the deletion of 2bp (allele 2), in exon 17, using Zinc Finger Nuclease editing (XCell Science). **B.** Sanger sequencing analysis of the genomic fragment around the two mutation sites, from control and KO iPSCs. The red rectangle shapes confirm the 2bp deletion (left) and the 4bp insertion (right), in the KO iPSC line. **C.** Quantitative RT-qPCR analysis of *CNTNAP2* expression in control and KO iPSCs, showing significantly decreased *CNTNAP2 mRNA* expression (n=2 biological replicates/genotype and 2-4 technical replicates/genotype. **D.** Western blot analysis that shows CNTNAP2 expression in D30 control organoids, and absence in D30 KO organoids. Each lane corresponds to one biological replicate. **E.** Karyotype analysis of control and KO iPSCs that confirms the normal karyotype of both cell lines. **F.** Quantitative RT-qPCR analysis (left) of the pluripotency markers NANOG and OCT-3/4 expression, and representative fluorescent images (right) from control and KO iPC, confirming the pluripotency of both cell lines. Data points represent different biological replicates collected at different time points. **G.** Representative fluorescent images from D30 control and KO organoids showing the expression pattern of the NPC marker SOX2 and the neuronal marker MAP2.

**Supplementary Fig. 2**

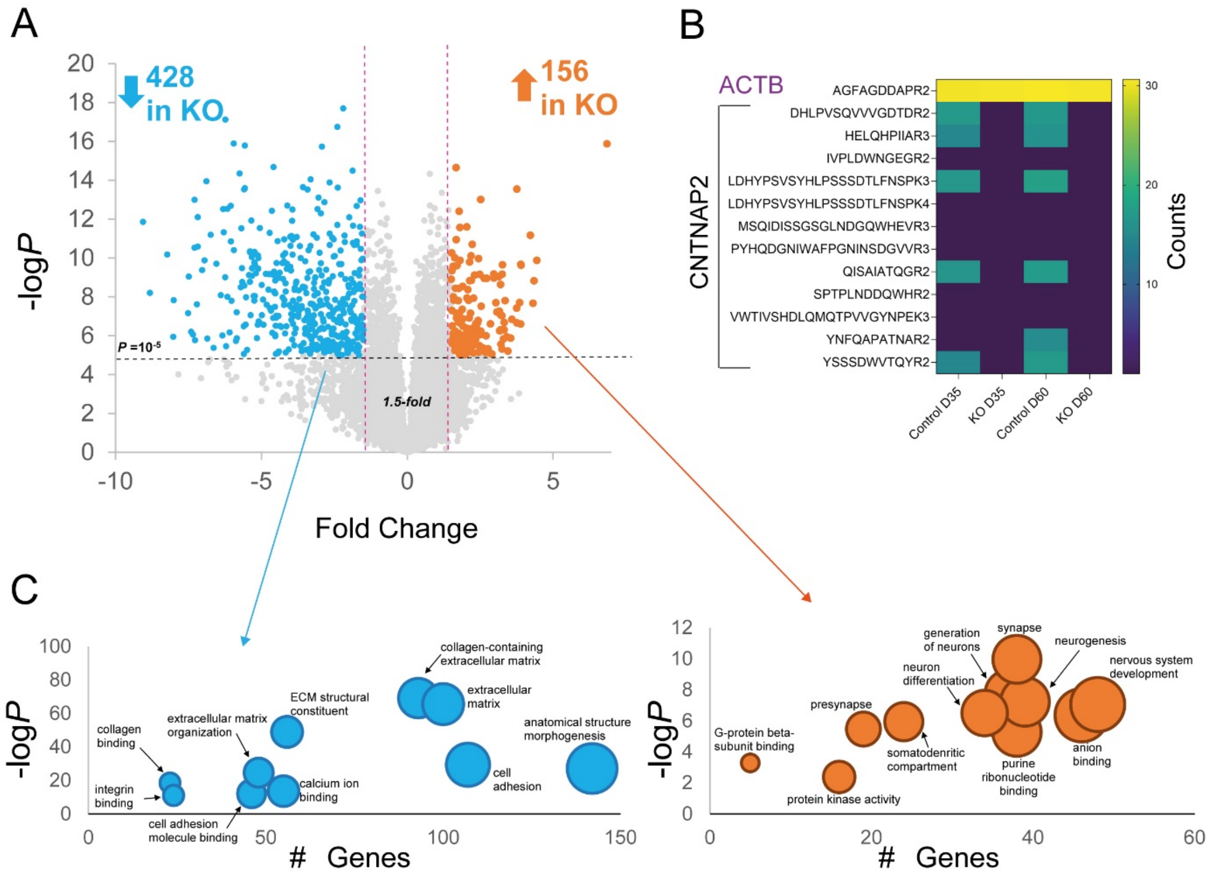


**Supplementary Figure 2 Additional Proteomics data A.** Volcano plot of D60 organoids proteomics experiment highlighting upregulated and downregulated peptides in KO samples. X-axis demonstrates the log-transformed fold change in abundance (KO/control) and the Y-axis indicates the log-transformed P values associated with individual peptides. A cut-off of ±1.5 fold-change (dashed vertical lines) and P value >10-5 (dashed horizontal line) was applied for differentially expressed peptides. **B.** Heatmap of CNTNAP2 and ACTB peptide counts in D30 and D60 proteomics (KO or control). Numbers in peptide names indicate peptide charge. **C.** GO analysis with g:Profiler for downregulated (left) and upregulated (right) peptides in KO. The Y-axis indicates the log-transformed P values; statistical analysis using g:GOSt (Fisher's one-tailed test).

See also Fig. 2 and Sup. Tables 3, 4, 7

**Supplementary Fig. 3**

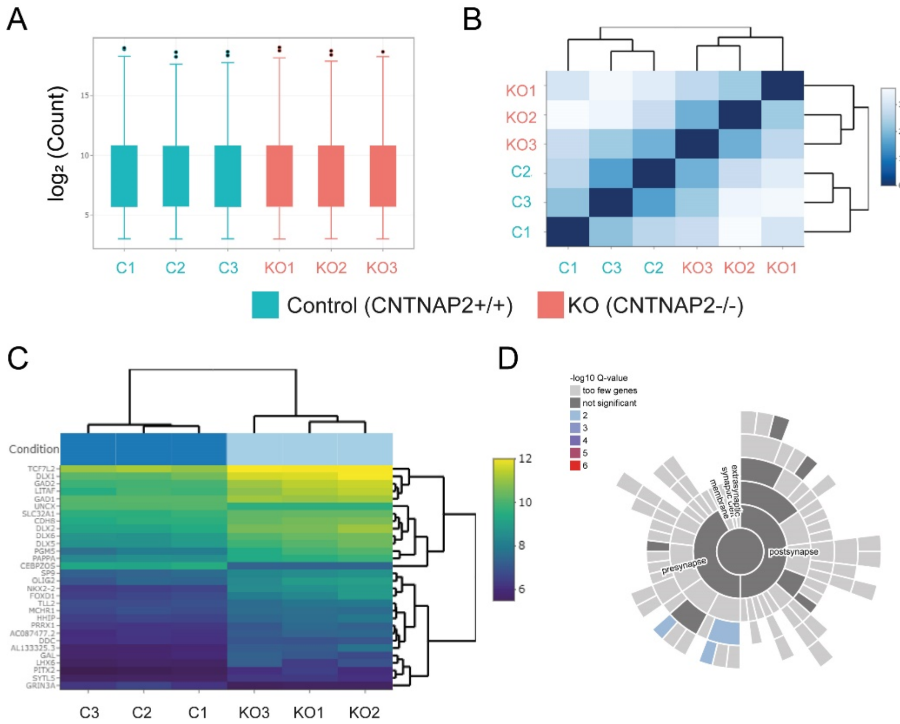


**Supplementary Figure 3 Additional RNAseq data A.** Normalisation of original RNAseq counts to adjust for various factors such as variations of sequencing yield between samples. These normalised read counts were used to accurately determine differentially expressed genes. **B.** Heatmap of overall similarity among samples assessed by the euclidean distance between samples. This method was used to examine which samples are similar/different to each other. The shorter the distance, the more closely related the samples are. **C.** Bi-clustering heatmap visualising the expression profile of the top 30 DEG sorted by their adjusted *P*-value by plotting their $log_2$ transformed expression values in samples. This analysis is useful to identify co-regulated genes across the experimental groups. **D.** SYNGO GO analysis of downregulated (left) and upregulated (right) DEG from KO RNAseq analysis of cerebral organoids.

See also Fig. 3 and Sup. Tables 5, 6, 7

### Supplementary Fig. 4

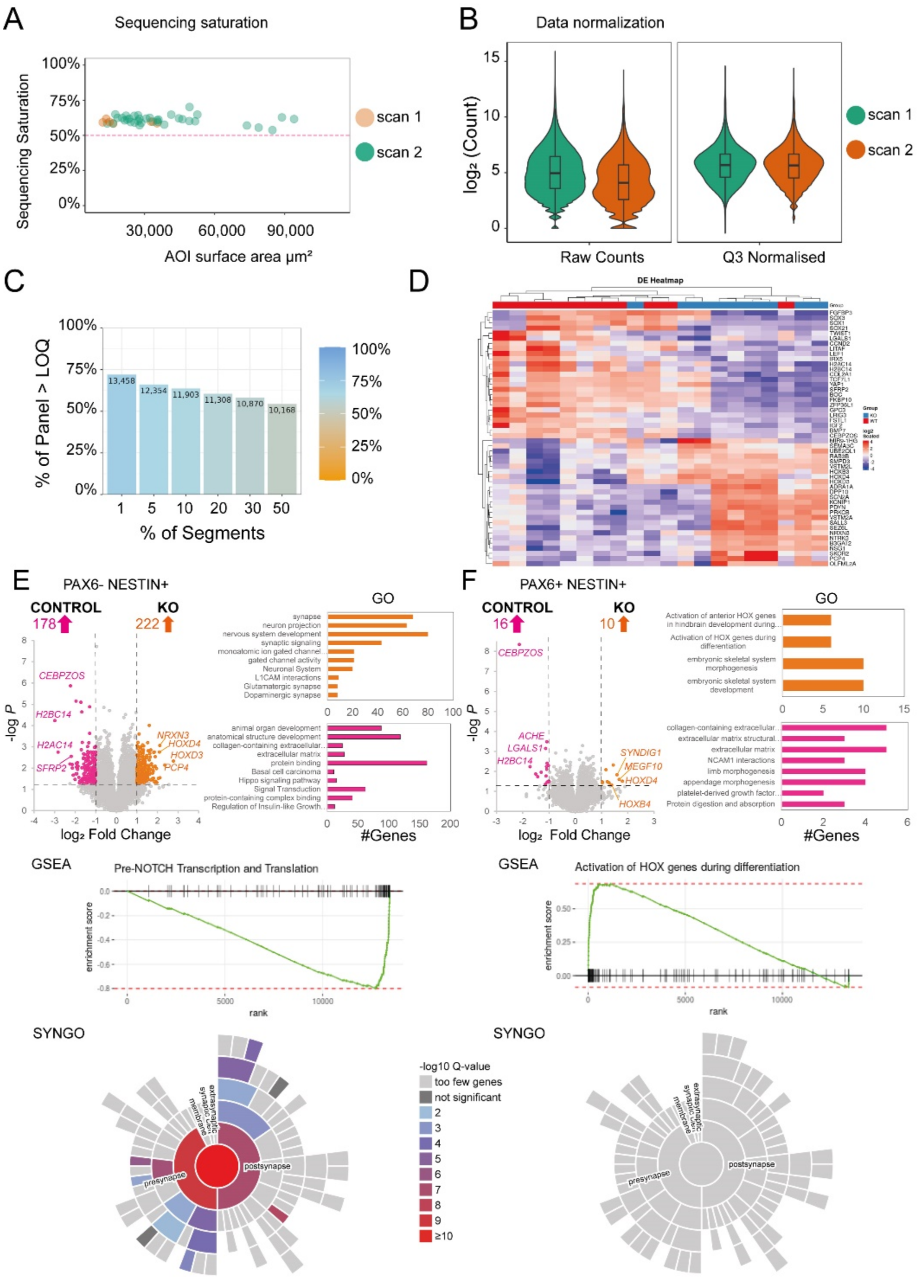


**Supplementary Figure 4 Additional spatial transcriptomics data. A.** Sequencing quality saturation shown per µm$^2$, ensuring sensitivity of low expressors. **B.** and **C.** Data was normalised to the third quartile (Q3) to account for differences in cellularity and AOI Size. We analysed 42 AOIs, 0 AOI were below the 50% warning, and we detected 18,676 total targets.13,457 genes normalised by 3rd quartile, expressed above Limit of Quantitation (LOQ) in at least 1% of AOIs. **D.** Heatmap of top 50 DEG sorted by genotype and $log_2$ fold change. **E.** and **F.** *Left:* Volcano plot of D30 spatial transcriptomics experiment highlighting upregulated and downregulated differentially expressed genes (DEG) in PAX6$^-$/NESTIN$^+$ or PAX6$^+$/NESTIN$^+$ in KO or control samples. X-axis demonstrates the $log_2$-transformed fold change in abundance in and the Y-axis indicates the negative log-transformed *P*adj (adjusted *P*) values associated with individual mRNAs. A cut-off of ±1 $log_2$ fold-change (dashed vertical lines) and log *P* value >1.3 (dashed horizontal line) was applied. *Right:* Gene ontology analysis of DEG with g:Profiler. Top categories (Biological Process, Molecular Function and Cellular Compartment) are shown for upregulated and downregulated DEG in the groups depicted. Statistical analysis was carried out using g:GOSt (Fisher's one-tailed test).

See also See also Fig. 4 and Sup. Tables 5, 6, 7
