## Supplementary Table 7 for "GABA/Glutamate neuron differentiation imbalance and increased AKT/mTOR signalling in CNTNAP2^-/-^ cerebral organoids"

**Supplementary Table 7** Statistical analysis details

| **Test** | **Mean± S.E.M** | **Standard Deviation (SD)** | **Significance** | **Normality-Shapiro Wilk** | **Parameter** | **N** | **Batches /genotype** | **Descriptive Statistics** | **Figure** |
| --- | --- | --- | --- | --- | --- | --- | --- | --- | --- |
| Mann Whitney test | Control: 1±0.037  KO: 0.907±0.029 | Control: 0.1652  KO: 0.127 | Control vs KO p=0.024 | Control; p=0.017  KO; p=0.105 | Normalised projected surface area, D30 organoids | Control (19)  KO (19) | 2 | Two-tailed, Mann Whitney U=103.5 | **Fig.1D** |
| Unpaired t test | Control: 1±0.035  KO: 1.031±0.057 | Control: 0.152  KO: 0.229 | Control vs KO p=0.633 | Control; p=0.76  KO; p=0.592 | Normalised projected surface area, D60 organoids | Control (19)  KO (16) | 2 | Two-tailed, t=0.4818, df=33 | **Fig.1D** |
| Unpaired t test | Control: 1±0.06  KO: 1.19±0.059 | Control: 0.17  KO: 0.156 | Control vs KO p=0.633 | Control; p=0.76  KO; p=0.043 | Normalised EdU/Ki67 | Control (8)  KO (7) | 3 | Two-tailed, t=2.238, df=13 | **Fig.1F** |
| Unpaired t test | Control: 0.399±0.079  KO: 0.733±0.018 | Control: 0.211  KO: 0.042 | Control vs KO p=0.005 | Control; p=0.28  KO; p=0.639 | Folds density % Area, D30 organoids | Control (7)  KO (5) | 2 | Two-tailed, Welch-corrected, t=4.058, df=6.653 | **Fig. 1H** |
| Unpaired t test | Control: 0.497±0.051  KO: 0.969±0.102 | Control: 0.154  KO: 0.288 | Control vs KO p=0.0007 | Control; p=0.309  KO; p=0.968 | Folds density % Area, D60 organoids | Control (9)  KO (8) | 2 | Two-tailed, t=4.267, df=15 | **Fig. 1H** |
| Unpaired t test | Control: 20345±1721  KO: 31699±3213 | Control: 4866  KO: 9088 | Control vs KO p=0.0076 | Control; p=0.154  KO; p=0.755 | VZ - Area | Control (8)  KO (8) | 4 | Two-tailed, t=3.115, df=14 | **Fig. 1J** |
| Unpaired t test | Control: 548.2±37.53  KO: 663.2±35.02 | Control: 112.6  KO: 99.05 | Control vs KO p=0.042 | Control; p=0.2358  KO; p=0.8079 | VZ - Perimeter | Control (9)  KO (8) | 4 | Two-tailed, t=2.223, df=15 | **Fig. 1J** |
| Unpaired t test | Control: 25.09±5.121  KO: 64.83±9.475 | Control: 15.36  KO: 26.8 | Control vs KO p=0.0017 | Control; p=0.934  KO; p=0.67 | %  Disorganised VZ | Control (9)  KO (8) | 4 | Two-tailed, t=3.809, df=15 | **Fig. 1J** |
| Unpaired t test | Control: 1±0.225  KO: 1.206±0.224 | Control: 0.748  KO: 0.809 | Control vs KO p=0.527 | Control; p=0.058  KO; p=0.082 | Normalised Phospho-S6 expression, D30 organoids | Control (11)  KO (13) | 4 | Two-tailed, t=0.642, df=22 | **Fig. 5C** |
| Unpaired t test | Control: 1±0.158  KO: 1.173±0.201 | Control: 0.549  KO: 0.699 | Control vs KO p=0.506 | Control; p=0.074  KO; p=0.256 | Normalised Phospho-AKT expression, D30 organoids | Control (12)  KO (12) | 4 | Two-tailed, t=0.676, df=22 | **Fig. 5C** |
| Unpaired t test | Control: 1±0.066  KO: 1.78±0.249 | Control: 0.229  KO: 0.864 | Control vs KO p=0.01 | Control; p=0.192  KO; p=0.193 | Normalised Phospho-S6 expression, D60 organoids | Control (12)  KO (12) | 3 | Two-tailed, Welch-corrected, t=3.023, df=12.54 | **Fig. 5C** |
| Mann Whitney test | Control: 1±0.08  KO: 1.297±0.086 | Control: 0.267  KO: 0.287 | Control vs KO p=0.023 | Control; p=0.039  KO; p=0.952 | Normalised Phospho-AKT expression, D60 organoids | Control (11)  KO (11) | 3 | Two-tailed, U=26 | **Fig. 5C** |
| Unpaired t test | Control: 42.58±4.229  KO: 41.87±3 | Control: 10.36  KO: 6.729 | Control vs KO p=0.898 | Control; p=0.974  KO; p=0.939 | %PAX6^+^ cell fraction | Control (6)  KO (5) | 3 | Two-tailed, t=0.131, df=9 | **Fig. 5D** |
| Unpaired t test | Control: 46.64±2.4  KO: 47.89±3.7 | Control: 7.228  KO: 11.72 | Control vs KO p=0.785 | Control; p=0.174  KO; p=0.28 | %SOX2^+^ cell fraction | Control (9)  KO (10) | 6 | Two-tailed, t=0.276, df=17 | **Fig. 5D** |
| Unpaired t test | Control: 5.926±0.581  KO: 8.977±0.556 | Control: 1.745  KO: 1.758 | Control vs KO p=0.0015 | Control; p=0.304  KO; p=0.793 | % MAP2^+^ Area | Control (9)  KO (10) | 6 | Two-tailed, t=3.79, df=17 | **Fig. 5D** |
| Mann Whitney test | Control: 0.699±0.245  KO: 3.373±0.791 | Control: 0.734  KO: 2.624 | Control vs KO p=0.016 | Control; p=0.0026  KO; p=0.306 | % GAD1+ Area | Control (9)  KO (11) | 4 | Two-tailed, Mann Whitney U=18 | **Fig. 5D** |
| Mann Whitney test | Control: 18.94±4.468  KO: 8.047±2.623 | Control: 13.40  KO: 8.295 | Control vs KO p=0.0435 | Control; p=0.1987  KO; p=0.02 | %TBR1^+^ cell fraction | Control (9)  KO (10) | 4 | Two-tailed, Mann Whitney U=20 | **Fig. 5D** |
| Nested t test | Control: 1.019±0.0713  KO: 0.11±0.034 | Control: 0.142  KO: 0.068 | Control vs KO p<0.0001 | Control; p=0.5  KO; p=0.3 | Relative CNTNAP2 mRNA expression | Control (6)  KO (6) | 3 | Two-tailed, t=12.52, df=10,  F=156.7, DFn=1, Dfd=10 | **Sup Fig.1C** |
| One-way ANOVA with Dunnett’s post hoc | Control: 17.69±4.8  KO: 14.42±2.854  Human brain: 1.007±0.004 | Control: 12.7  KO: 7.55  Human brain:0.011 | Control vs KO p=0.911  Control vs Human brain p=0.035  KO vs Human brain  p=0.009 | Control; p=0.842  KO; p=0.957  Human brain; p=0.0002 | Relative NANOG mRNA expression | Control (7)  KO (7)  Human brain (7) | 3 | Brown-Forsythe ANOVA test F*(DFn, DFd)=7.516 (2.000, 9.770), p=0.0106  Welch’s ANOVA test W(DFn, DFd)=15.77(2.000, 8.000) p=0.0017 | **Sup Fig.1F** |
| One-way ANOVA with Dunnett’s post hoc | Control: 24.5±2.56  KO: 22.74±1.76  Human brain: 1.015±0.01 | Control: 6.78  KO: 4.66  Human brain:0.027 | Control vs KO p=0.92  Control vs Human brain p=0.0003  KO vs Human brain  p<0.0001 | Control; p=0.657  KO; p=0.887  Human brain; p<0.0001 | Relative OCT-3/4 mRNA expression | Control (7)  KO (7)  Human brain (7) | 3 | Brown-Forsythe ANOVA test F*(DFn, DFd)=52.99 (2.000, 10.64), p<0.0001  Welch’s ANOVA test W(DFn, DFd)=108.7 (2.000, 8.000) p<0.0001 | **Sup Fig.1F** |
