## Supplementary Table 8 for "GABA/Glutamate neuron differentiation imbalance and increased AKT/mTOR signalling in CNTNAP2^-/-^ cerebral organoids"

**Supplementary Table 8, List of Antibodies**

| **Antibody** | **Vendor** | **Catalog** | **Species** | **Dilution** |
| --- | --- | --- | --- | --- |
| Antibody | Vendor | Catalog | Species | Dilution |
| Anti-AKT | Cell Signaling Technologies | 9272S | Rabbit | 1:1000 |
| Anti -Caspr/CNTNAP2 | Abcam | ab153856 | Rabbit | 1:2000 |
| Anti-GAD1 | Sigma-Aldrich | MAB5406 | Rabbit | 1:200 |
| Anti-HSC70 | Santa Cruz Biotechnology | sc-7298 | Mouse | 1:1000 |
| Anti-Ki67 | Agilent Technologies | M724029-2 | Mouse | 1:200 |
| Anti-NANOG | Cell Signaling Technologies | 3580S | Rabbit | 1:100 |
| Anti-NESTIN | Cell Signaling Technologies | 33475 | Mouse | 1:200 |
| Anti-OCT-3/4 | Santa Cruz Biotechnology | sc-5279 | Mouse | 1:100 |
| Anti-phospho-AKT (S473) | Cell Signaling Technologies | 9271S | Rabbit | 1:1000 |
| Anti-PAX6 | Cell Signaling Technologies | 60433S | Rabbit | 1:200 |
| Anti-phospho-rpS6 (S240/244) | Cell Signaling Technologies | 2211S | Rabbit | 1:1000 |
| Anti-rpS6 | Santa Cruz Biotechnology | sc74459 | Mouse | 1:1000 |
| Anti-SOX2 | Santa Cruz Biotechnology | sc365823 | Mouse | 1:200 |
| Anti-TBR1 | Cell Signaling Technologies | 49661S | Rabbit | 1:250 |
| Anti-TUJ1 | Santa Cruz Biotechnology | sc-80005 | Mouse | 1:200 |
| Donkey-anti mouse 680 | LI-COR Biosciences | 926-68072 | Mouse | 1:1000 |
| Donkey-anti rabbit 800 | LI-COR Biosciences | 926-32213 | Rabbit | 1:1000 |
| Goat-anti mouse 800 | LI-COR Biosciences | 926-32210 | Mouse | 1:1000 |
| Goat-anti rabbit 680 | LI-COR Biosciences | 926-68071 | Rabbit | 1:1000 |
| Goat-anti-mouse 488 | Thermo Fisher Scientific | A-11001 | Mouse | 1:1000 |
| Goat-anti-mouse 568 | Thermo Fisher Scientific | A-11004 | Mouse | 1:1000 |
| Goat-anti-rabbit 488 | Thermo Fisher Scientific | A-11008 | Rabbit | 1:1000 |
| Goat-anti-rabbit 568 | Thermo Fisher Scientific | A-11011 | Rabbit | 1:1000 |
