## Supplementary Figure 5 for "GABA/Glutamate neuron differentiation imbalance and increased AKT/mTOR signalling in CNTNAP2^-/-^ cerebral organoids"

### Western Blots

Membranes – Figure 5

D30 cerebral organoids

Phospho - S6 (240/244)

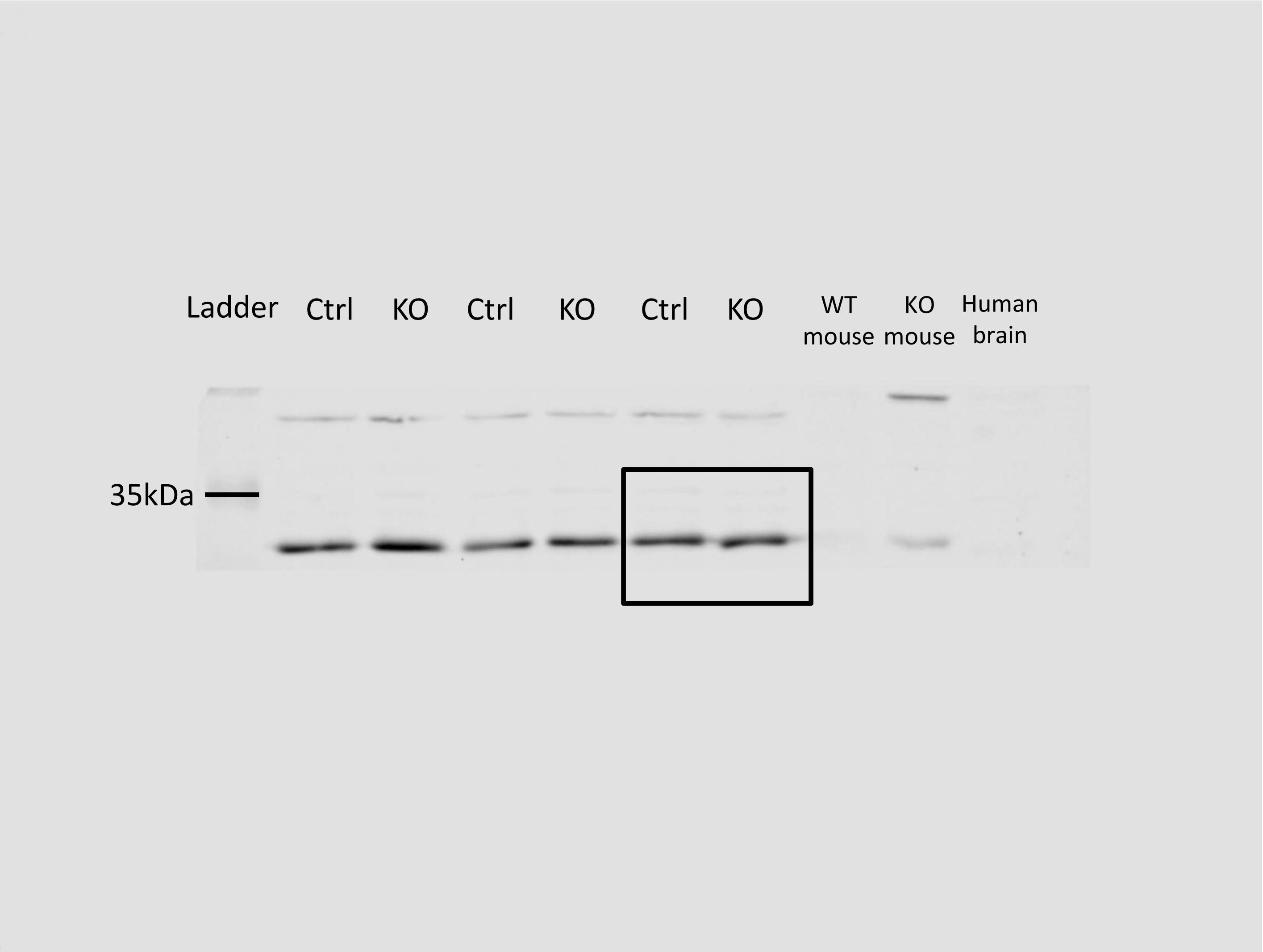

D30 cerebral organoids

Total S6

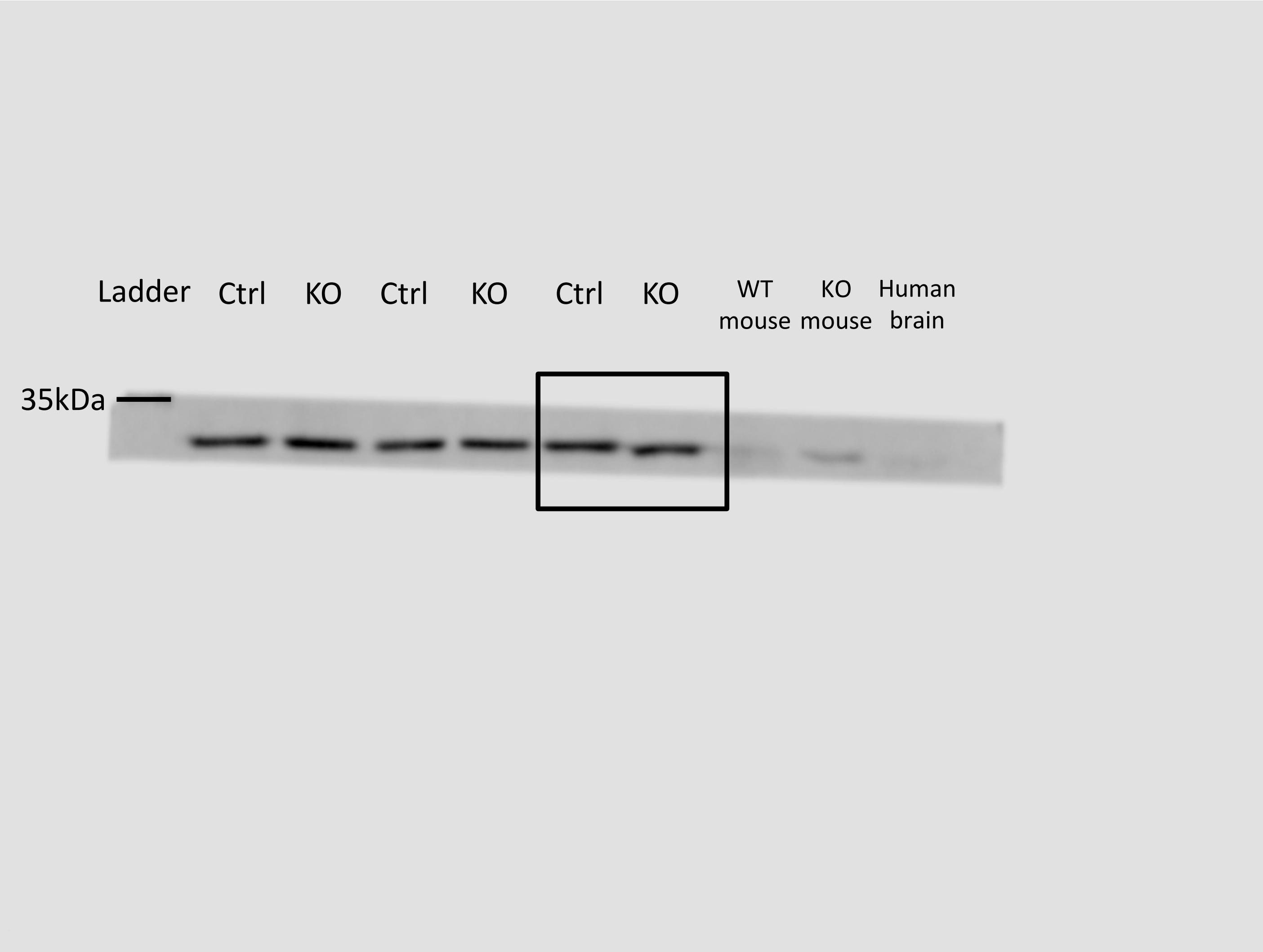

D30 cerebral organoids

HSC70

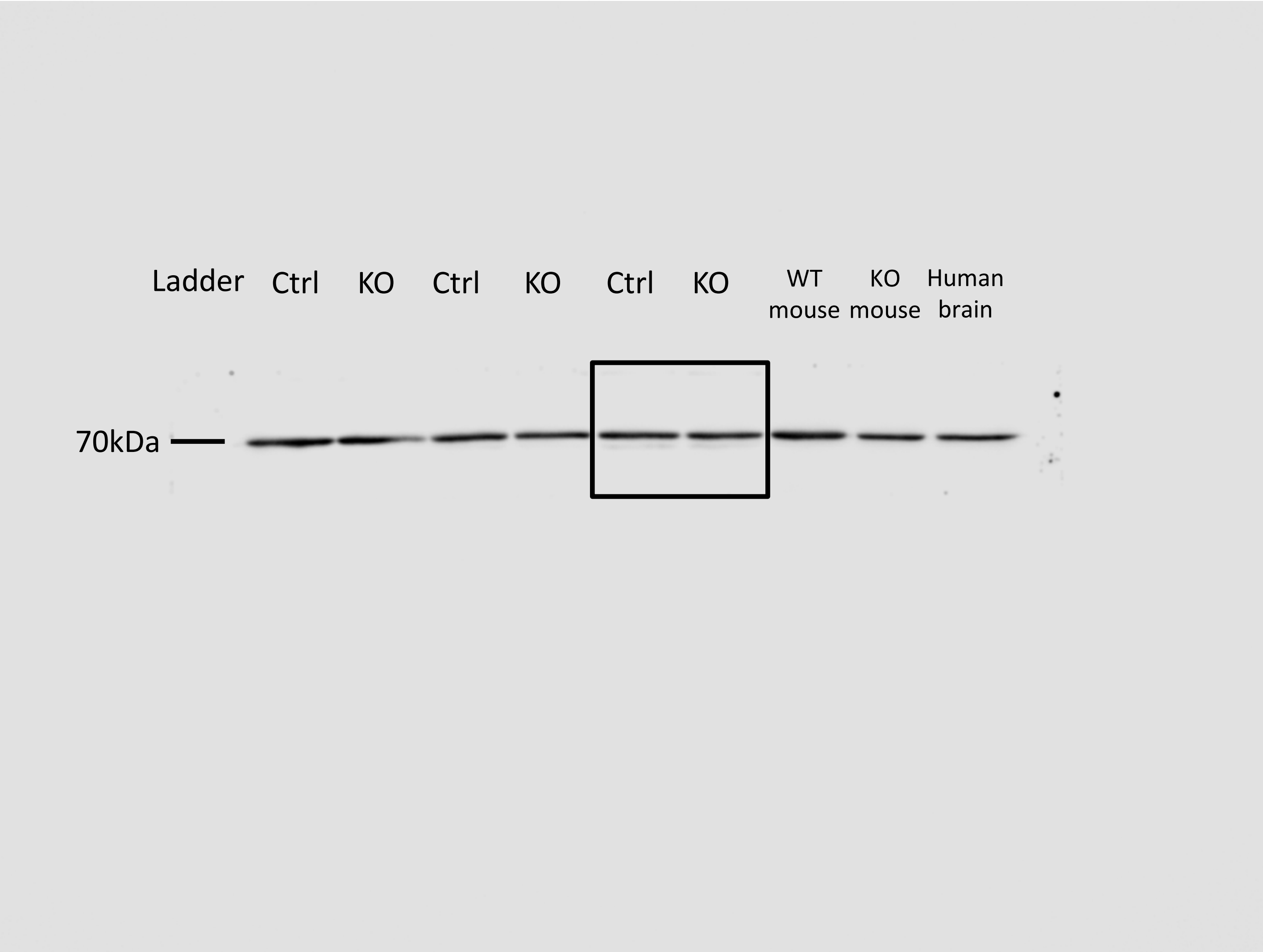

### D30 cerebral organoids

Phospho –AKT (S473)

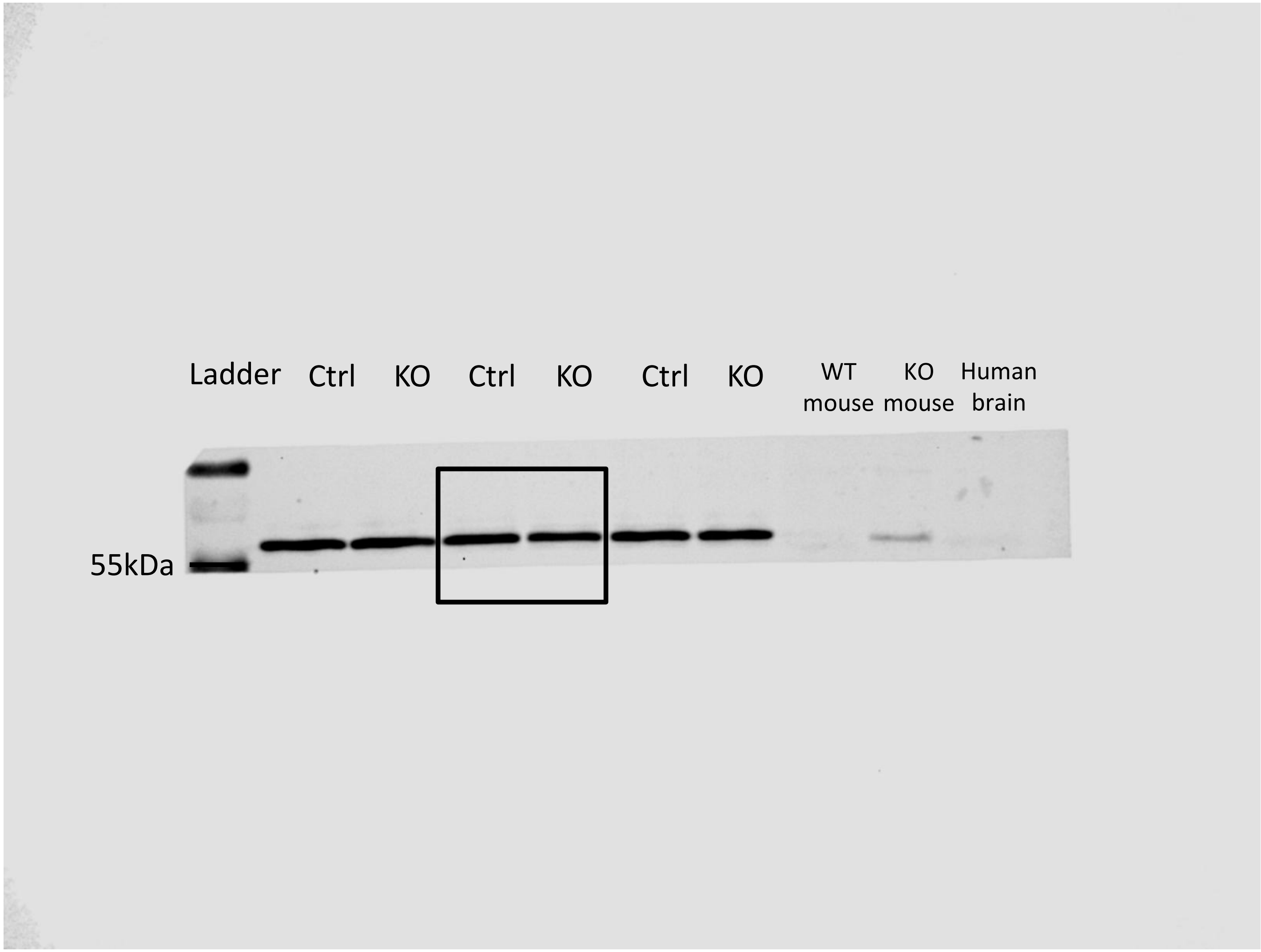

D30 cerebral organoids

Total AKT

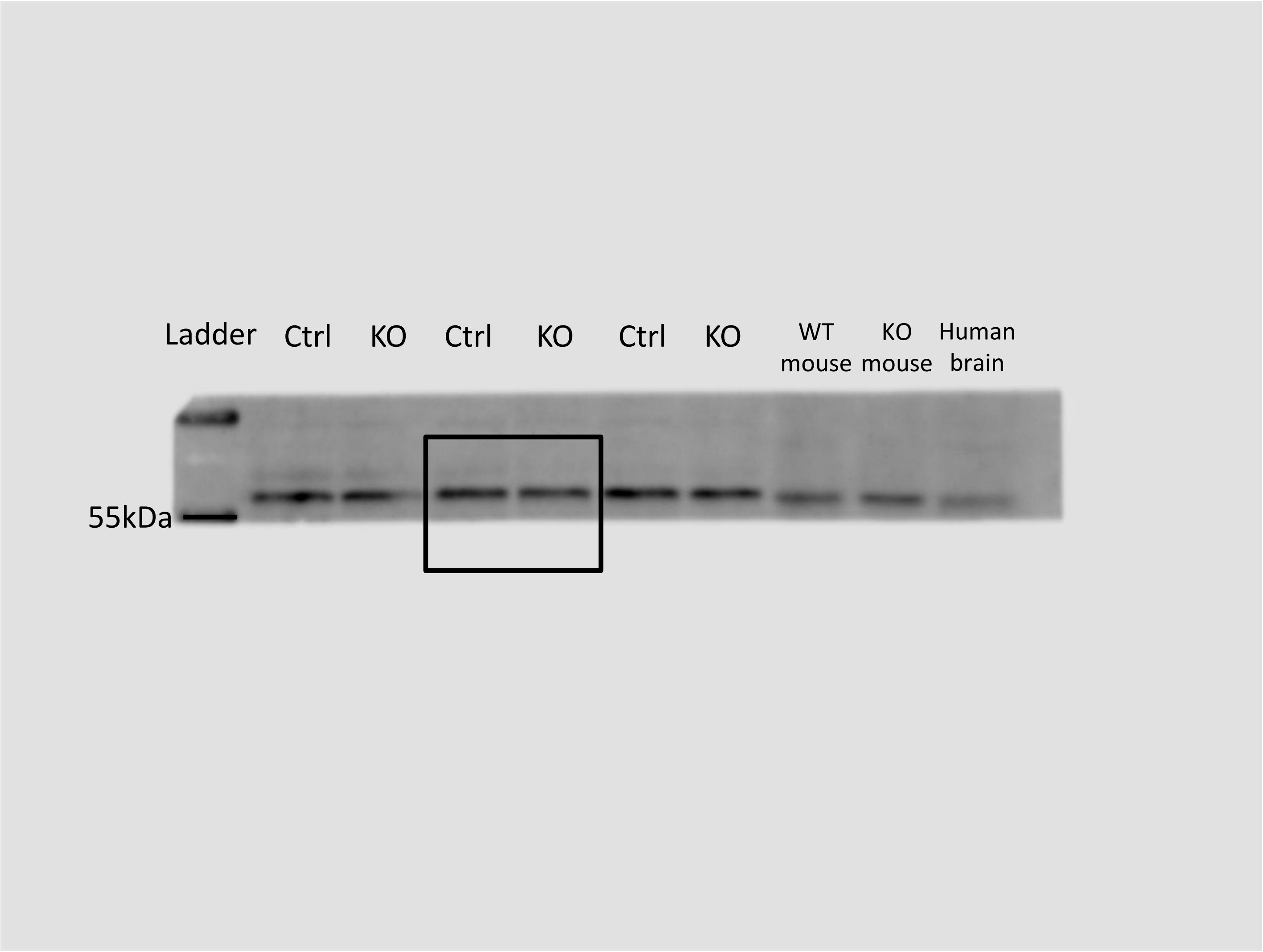

D30 cerebral organoids

HSC70

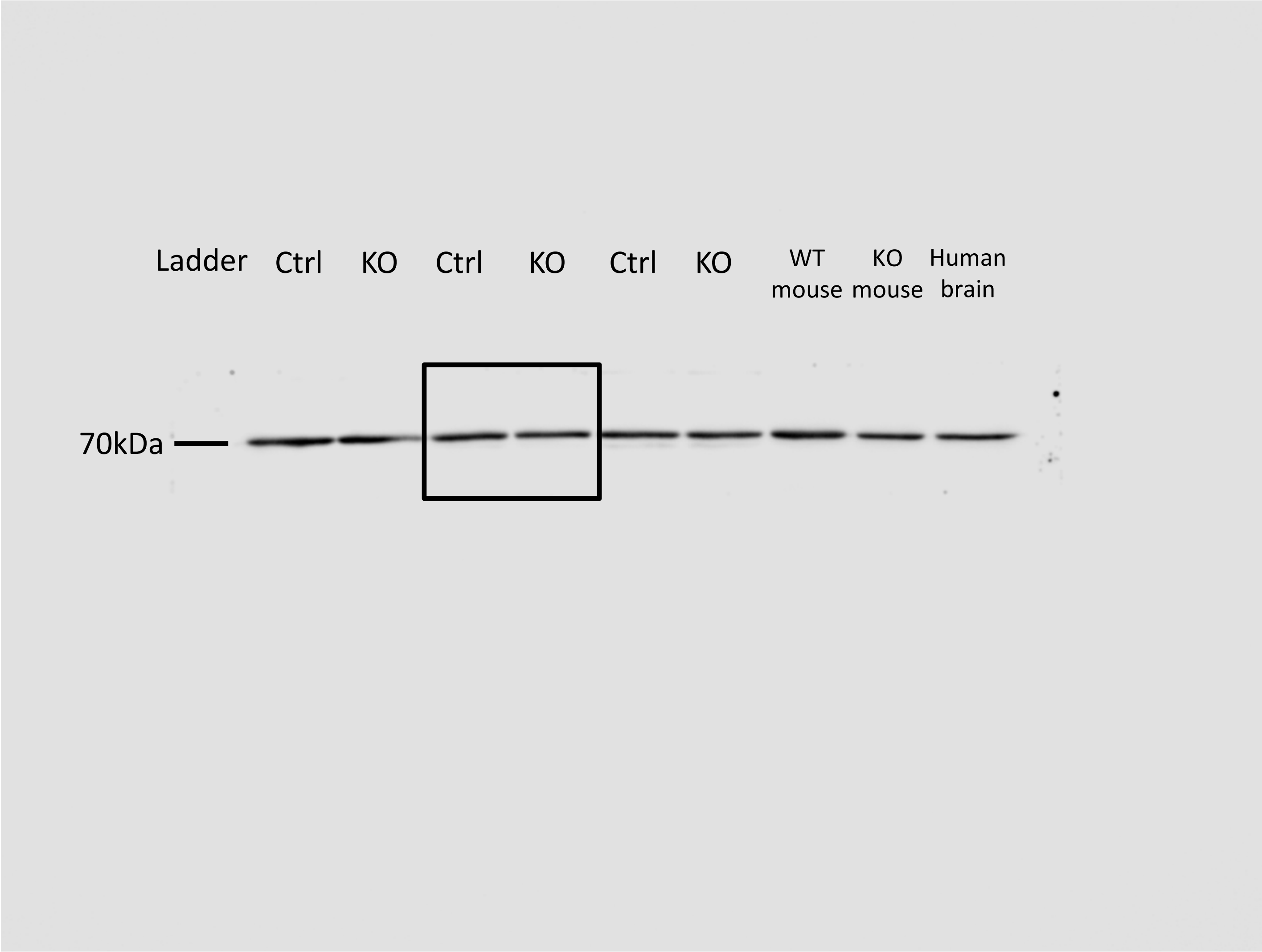

D60 cerebral organoids

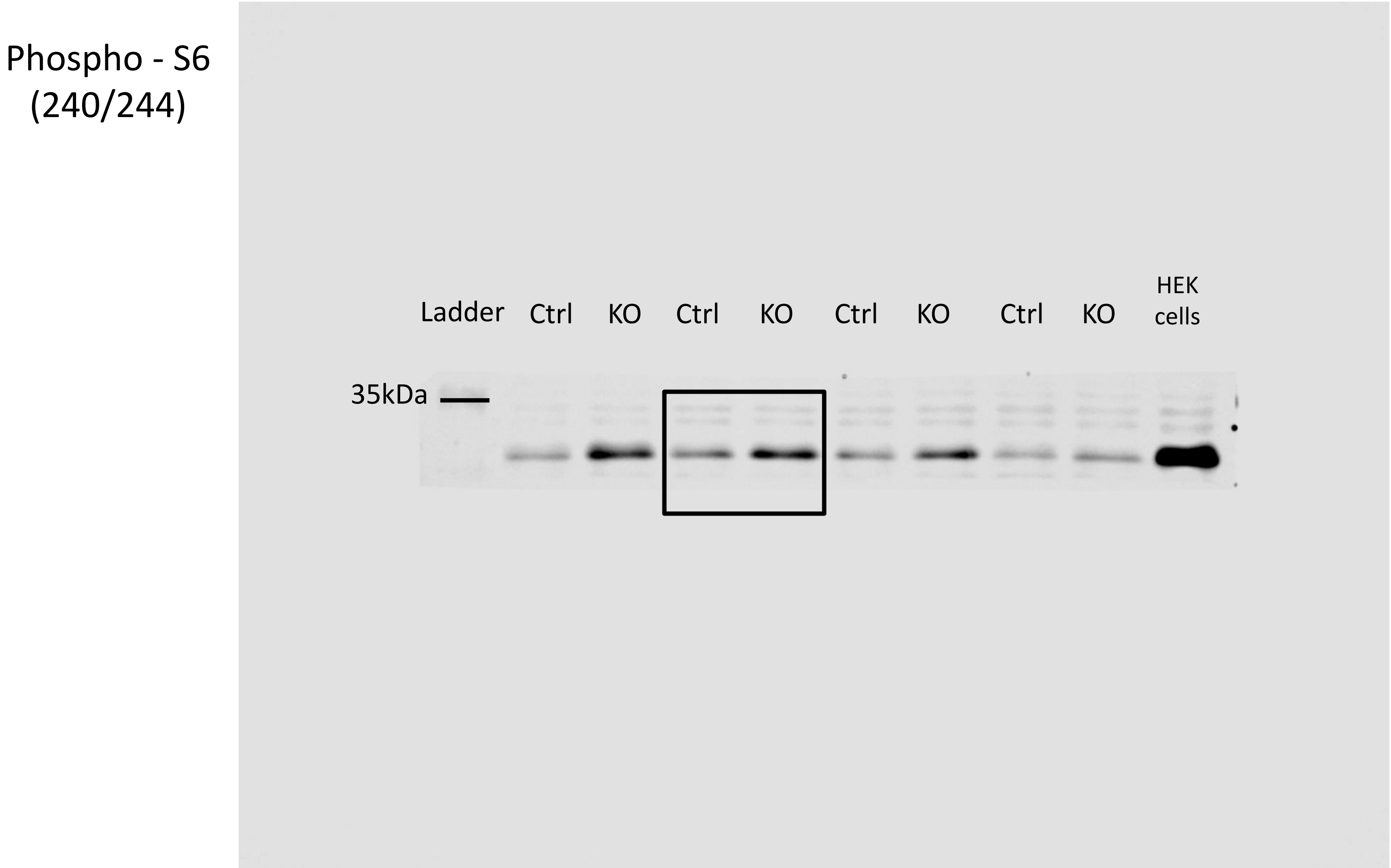

D60 cerebral organoids

Total S6

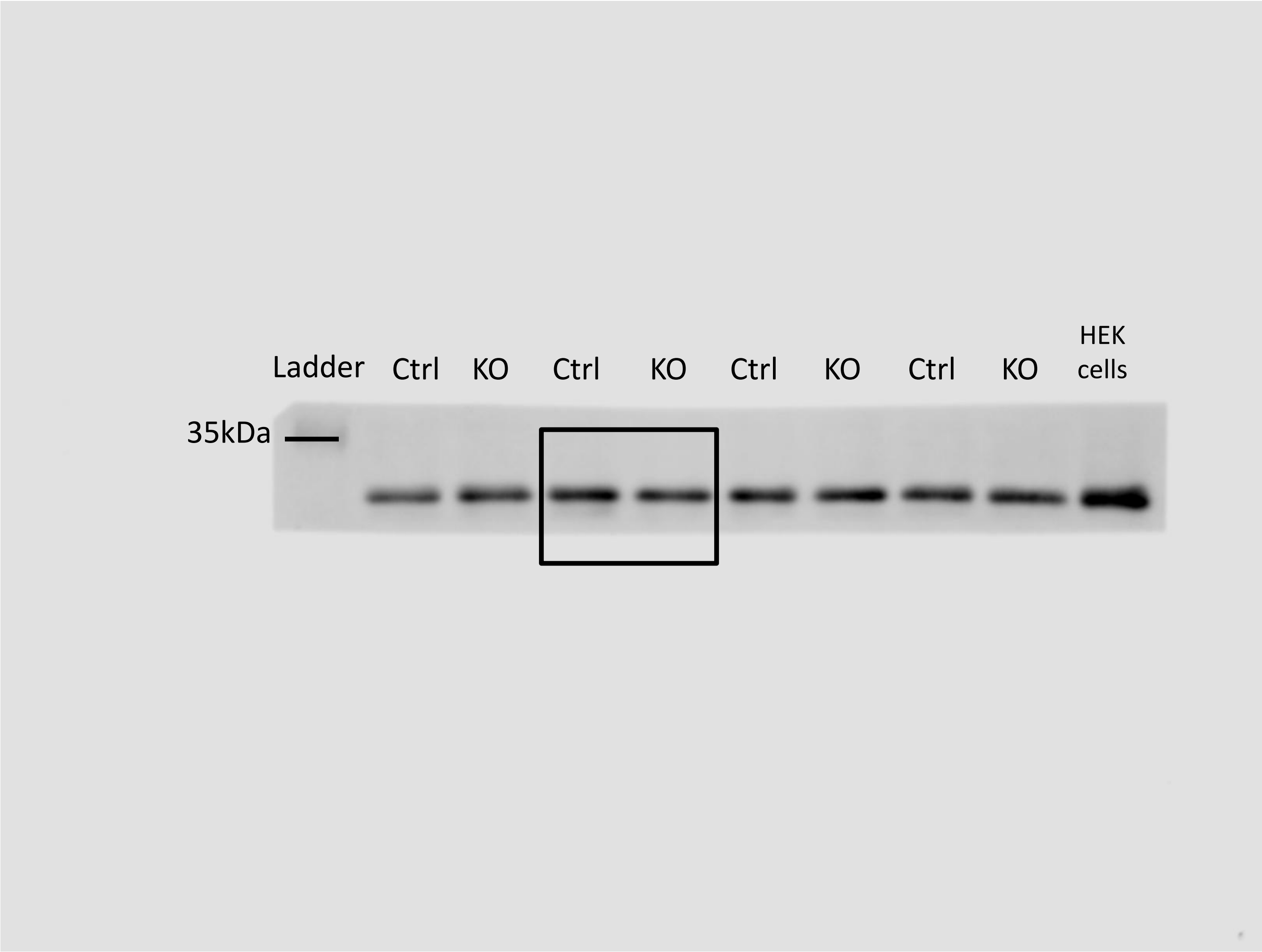

D60 cerebral organoids

HSC70

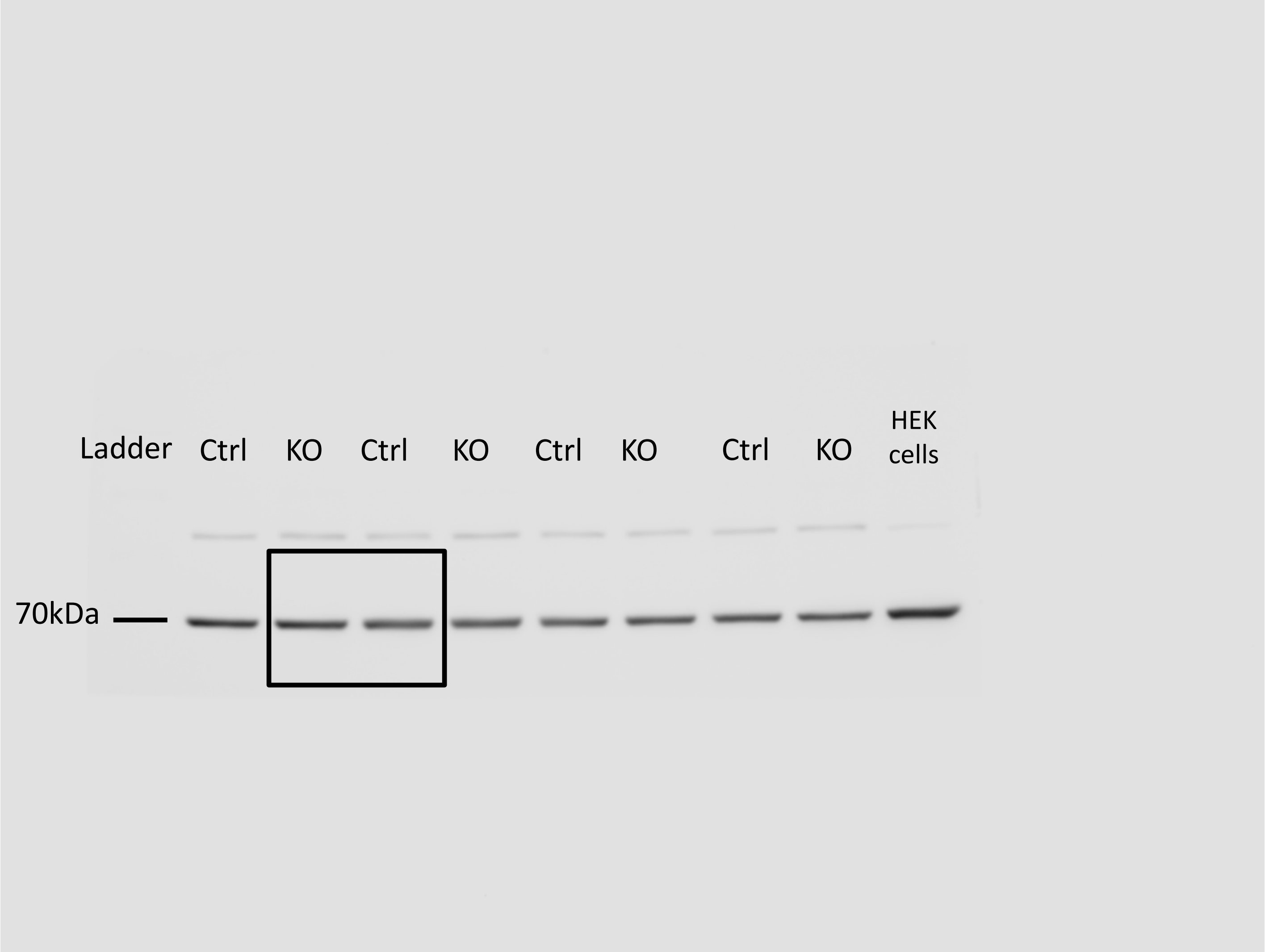

D60 cerebral organoids

Phospho –AKT (S473)

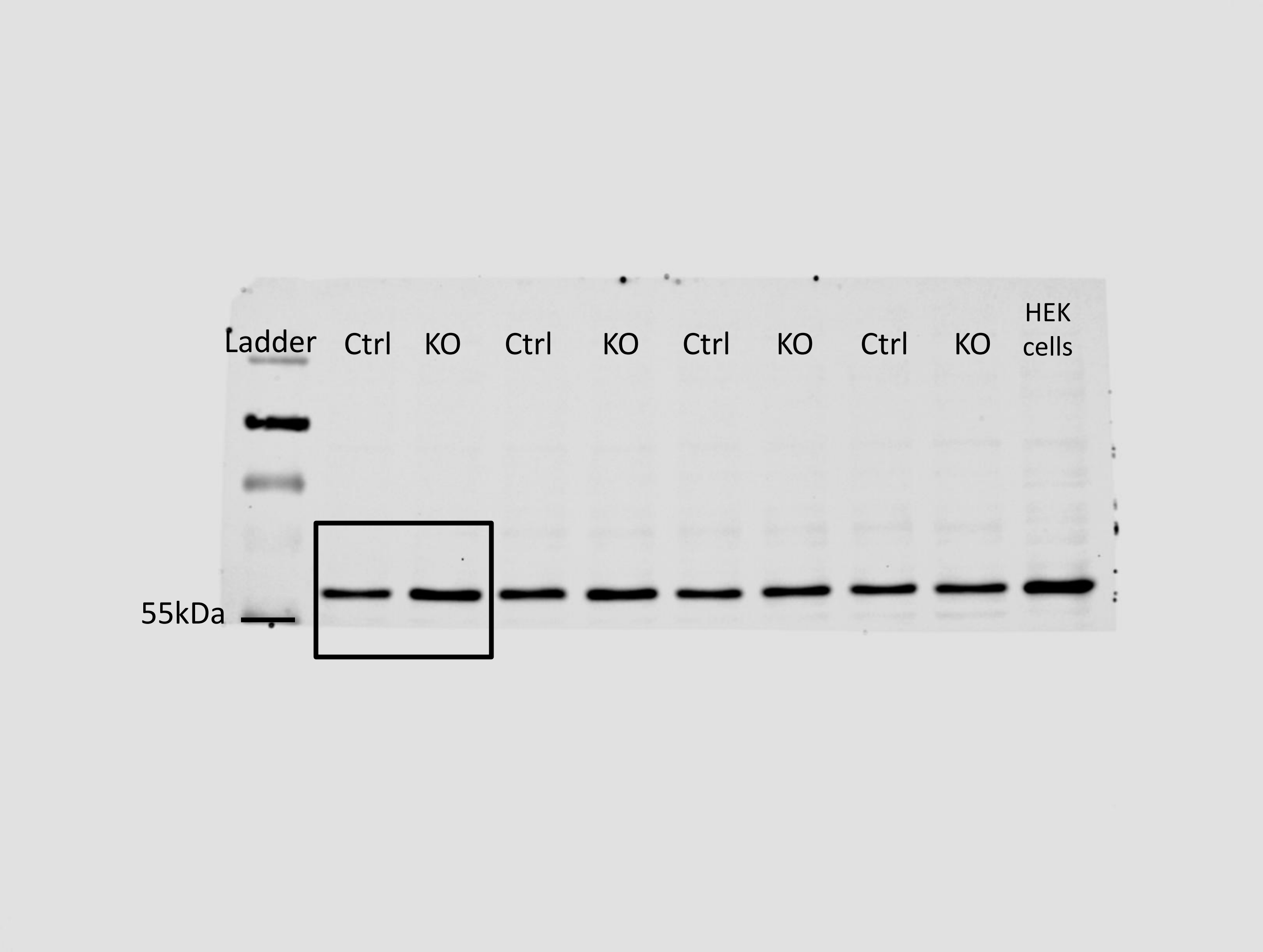

D60 cerebral organoids

Total AKT

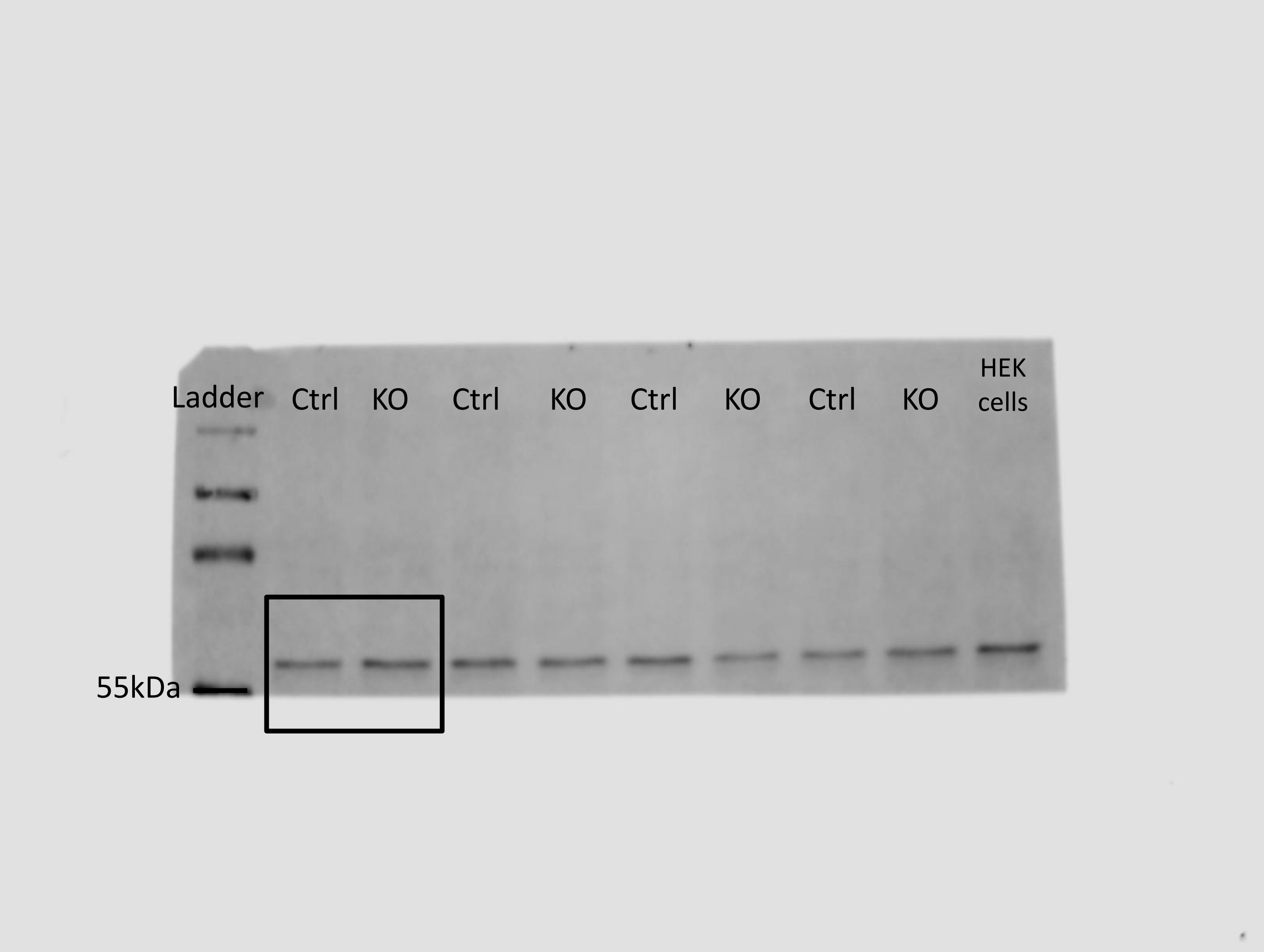

D60 cerebral organoids

HSC70

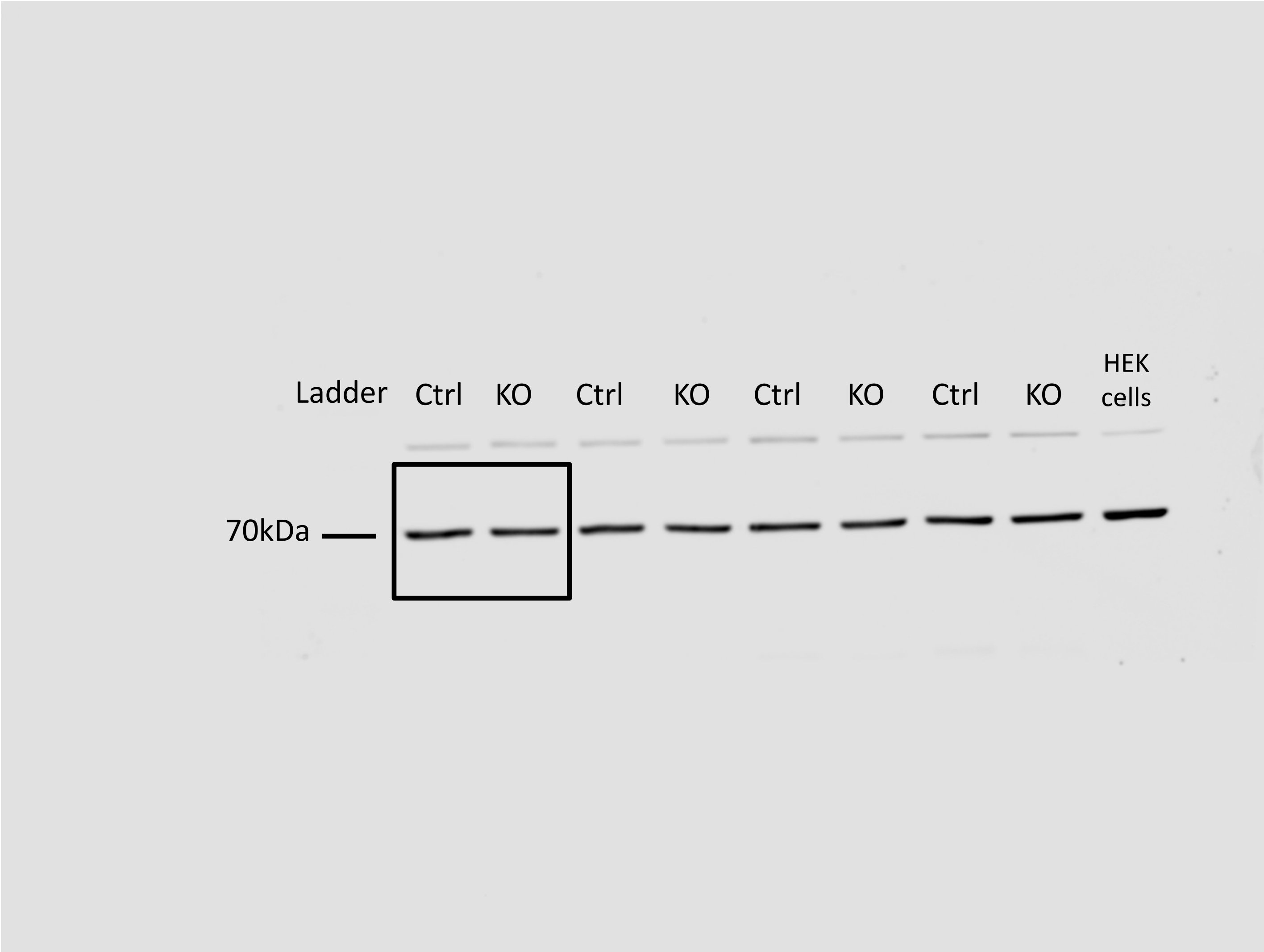
